# Circuit and cell-specific contributions to decision making involving risk of explicit punishment in male and female rats

**DOI:** 10.1101/2023.01.15.524142

**Authors:** Leah M. Truckenbrod, Sara M. Betzhold, Alexa-Rae Wheeler, John Shallcross, Sarthak Singhal, Scott Harden, Marek Schwendt, Charles J. Frazier, Jennifer L. Bizon, Barry Setlow, Caitlin A. Orsini

**Author notes:** Corresponding author: Correspondance should be addressed to Dr. Caitlin A. Orsini, Department of Psychology & Neurology, The University of Texas at Austin, 1601B Trinity Street, Austin, TX 78712.

## Abstract

Decision making is a complex cognitive process that recruits a distributed network of brain regions, including the basolateral amygdala (BLA) and nucleus accumbens shell (NAcSh). Recent work suggests that communication between these structures, as well as activity of cells expressing dopamine D2 receptors (D2R) in the NAcSh, are necessary for some forms of decision making; however, the contributions of this circuit and cell population during decision making under risk of punishment are unknown. The current experiments addressed this question using circuit- and cell type-specific optogenetic approaches in rats during a decision-making task involving risk of punishment. In Experiment 1, Long-Evans rats received intra-BLA injections of halorhodopsin or mCherry (control) and in Experiment 2, D2-Cre transgenic rats received intra-NAcSh injections of Cre-dependent halorhodopsin or mCherry. Optic fibers were implanted in the NAcSh in both experiments. Following training in the decision-making task, BLA→NAcSh or D2R-expressing neurons were optogenetically inhibited during different phases of the decision process. Inhibition of the BLA→NAcSh during deliberation (the time between trial initiation and choice) increased choice of the large, risky reward (increased risk taking). Similarly, inhibition during delivery of the large, punished reward increased risk taking, but only in males. Inhibition of D2R-expressing neurons in the NAcSh during deliberation increased risk taking. In contrast, inhibition of these neurons during delivery of the small, safe reward decreased risk taking. These findings extend our knowledge of the neural dynamics of risk taking, revealing sex-dependent circuit recruitment and dissociable activity of selective cell populations during decision making.

## Introduction

Decision making consists of multiple cognitive operations that occur in synchrony with one another to give rise to choice behavior (Fobbs and Mizumori, 2017; Orsini et al., 2019). For example, when faced with a decision, an individual must evaluate the available options and weigh their benefits against their costs (“deliberation”). Simultaneously, feedback from past decisions of a similar nature must be considered and integrated into this deliberative process. Thus, what appears to be a homogenous process is actually multifaceted and requires significant cognitive resources. The architecture of a decision is particularly important to understand in the context of neuropsychiatric conditions associated with impaired decision making. For instance, individuals with substance use disorder display increased risk-taking behavior (Chen et al., 2020), which is thought to be related to diminished sensitivity to risks of adverse consequences (Bechara and Damasio, 2002; Lejuez et al., 2005; Myers et al., 2017). This deficit suggests that the ability to use negative feedback to guide future decisions, rather than the deliberative process, may be compromised in these individuals.

These distinct components of decision making are mediated by temporally-specific recruitment of regions within the mesocorticolimbic circuit (Orsini et al., 2019). For example, activity within the basolateral amygdala (BLA) contributes to decision making involving risk of punishment differentially across the decision process (Orsini et al., 2017). Whereas activity of BLA glutamatergic neurons during deliberation biases choices toward larger, riskier rewards, activity of these neurons during evaluation of punished rewards guides future choice away from larger, riskier rewards. Activity in the BLA may influence risk taking in such a dissociable manner through divergent projections to other brain regions. For instance, BLA projections to prefrontal cortical areas may be important for deliberative processes of risk-based decision making, while BLA projections to the shell subregion of the nucleus accumbens (NAcSh; BLA→NAcSh) may be necessary for evaluation of punished rewards. Support for the latter possibility comes from work showing that disconnection of the BLA and NAcSh decreases the number of attempts to avoid a footshock in an active avoidance task (Ramirez et al., 2015). Although previous studies have shown a temporally-specific role for the BLA→NAcSh in decision making involving risk of reward omission (Bercovici et al., 2018), the temporal dynamics of this pathway in decision making involving risk of punishment remains unknown. This is an important consideration as the neurobiological mechanisms of risk-based decision making differ as a function of the type of risk involved (Orsini et al., 2015b).

Within the NAcSh, dopamine (DA) neurotransmission influences risk-based decision making by binding to DA receptors (Orsini et al., 2015b; Winstanley and Floresco, 2016). Unlike other forms of risk-based decision making in which both DA D1 and D2 receptors mediate aspects of choice behavior (St Onge and Floresco, 2009; St Onge et al., 2010), decision making involving risk of punishment selectively involves D2 receptors (D2Rs); systemic or intra-NAcSh administration of D2R agonists decreases choice of large, risky rewards, whereas administration of D1R ligands has no effect on choice performance (Simon et al., 2011; Mitchell et al., 2014). The use of pharmacological methods, however, precludes determination of the specific components of the decision process for which these receptors, and the cells on which they reside, are necessary. Like with BLA glutamatergic neurons, the activity of NAcSh D2R-expressing neurons may in fact subserve distinct aspects of decision making. Indeed, a recent study demonstrated that NAc D2R-expressing neurons selectively encode loss sensitivity during deliberation, but not outcome evaluation, in a decision-making task involving risk of reward omission (Zalocusky et al., 2016). Hence, engagement of NAcSh D2R-expressing neurons may be similarly selective to deliberation in decision making involving risk of punishment.

The current set of experiments were designed to evaluate the roles of the BLA→NAcSh and NAcSh D2R-expressing neurons during distinct components of decision making involving risk of punishment. Collectively, the results of these experiments advance our understanding of the neural dynamics underlying decision making.

## Materials and Methods

### Subjects

For experiments involving manipulations of the BLA→NAcSh, Long-Evans rats were obtained from Charles River (UF: Kingston, NY; UT Austin: Hollister, CA) at postnatal (P) day 50 and individually housed upon arrival. For experiments involving manipulations of D2R-expressing neurons in the NAcSh, male and female hemizygous D2-Cre rats [LE-Tg(Drd2-iCre)1Ottc] were procured from the Optogenetic Transgenic Technology Core (NIDA) through the Rat Research and Resource Center (RRRC#768, Missouri, USA). Hemizygous male and female rats were initially paired to begin the breeding colony, but after the colony was established, hemizygous male rats were mated with wild-type Long-Evans female rats obtained from Charles River (P90). Upon weaning (∼P21), hemizygous D2-Cre male and female rats were housed in groups of 3-4 (separately for each sex) until P60, at which point they were individually housed and underwent surgery. Experiments were conducted at both the University of Florida and The University of Texas at Austin, but there were no differences in baseline behavioral performance or in the effects of manipulations; consequently, datasets were merged and analyzed together. All rats were maintained on a reverse light cycle (lights on at 1900h), housed in ventilated cages with Sanichip bedding and had *ad libitum* access to water. During initial behavioral testing, rats were food-restricted to 85% of their free-feeding weight, with weights adjusted upward by 5 grams per week to account for growth. Once fully grown (∼250g for females, ∼350g for males), rats were fed 10 grams of food (soy-free; Envigo Teklad Irradiated Global 19% Protein Extruded Rodent Diet, #2919) each day after behavioral testing. Environmental enrichment in the form of nylabones were provided in each cage and replaced as needed. All procedures were conducted in accordance with the regulations and policies stipulated by the Institutional Animal Care and Use Committees at UF and UT Austin and adhered to ethical guidelines set forth by the National Institutes of Health.

### Apparatus

Behavioral testing was conducted in standard operant chambers (Coulbourn Instruments; 8 chambers, UF; 6 chambers, UT Austin), each of which was housed in sound-attenuating cabinets. A centrally located food trough projected 3 cm out from the front wall of each chamber and was equipped with photobeams to enable detection of entries into the trough. Each food trough was connected to a feeder, from which 45 mg soy-based food pellets (UF: Test Diet, 5TUL, Richmond, IN; UT Austin: Lab Supply, 5TUL, Houston, TX) were delivered into the trough. A nosepoke was positioned directly above the food trough and two retractable levers flanked each side of the food trough. The floor of each chamber consisted of stainless-steel rods through which scrambled footshocks were delivered via the floor’s connection to a shock generator (Coulbourn Instruments). A rotary joint (1 × 2, 200 µm core, Doric Lenses) was mounted on the ceiling of each operant chamber for use during optogenetic manipulations. Each chamber was interfaced with a computer running Graphic State 4.0, which controlled chamber hardware (e.g., lever extension, laser delivery) and concurrently recorded behavioral output (e.g., nosepokes, lever presses).

### Laser Delivery

For optogenetic inhibition of the BLA→NAcSh (Experiment 1), laser light (560 nm, 12-16 mW output, Shanghai Laser & Optics Century) was delivered bilaterally through optic fibers (200 µm core, 0.22 NA, Precision Fiber Products) implanted in the NAcSh of rats that received intra-BLA injections of halorhodopsin (eNpHR3.0 group) or mCherry (control group). For optogenetic inhibition of D2R-expressing neurons in the NAcSh (Experiment 2), laser light (560 nm, 8-10 mW output) was delivered bilaterally through optic fibers implanted in the NAcSh of D2-Cre rats that received intra-NAcSh injections of halorhodopsin (eNpHR3.0 group) or mCherry (control group). In all rats, light was transmitted from the laser to the rotatory joint through a patch cord (200 µm core, Thor Labs) and then from the rotary joint to the implanted optic fibers through two additional patch cords (200 µm core, 0.22 NA, Thor Labs) encased in stainless steel. The laser was interfaced with the computer running Graphic State 4.0 software to enable precise control of the timing of light delivery during specific task events.

### Surgical procedures

Rats were anesthetized with isoflurane gas (1-5% in O_2_) and were administered subcutaneous injections of buprenorphine (0.03 mg/kg), meloxicam (2 mg/kg) and 10 mL of 0.9% saline. Upon reaching a stable plane of anesthesia, rats’ hair on the superior aspect of the scalp was clipped, after which they were secured in the stereotax (Kopf Instruments). The scalp was disinfected with chlorohexidine and isopropyl alcohol swabs and ophthalmic ointment was applied to the eyes. A sterile adhesive surgical drape was then placed over the body.

#### Validation experiment surgeries

Prior to the behavioral manipulations in Experiment 2, RNAscope was used to verify that viral transduction of D2R-expressing neurons was selective to these neurons. In addition, *in vitro* electrophysiology experiments were conducted to confirm that optogenetic inhibition of D2R-expressing neurons had the intended physiological consequences. For rats in these validation experiments, an incision was made in the scalp and the skin was retracted with hemostats. After clearing the fascia, the skull was leveled to ensure that bregma and lambda were in the same horizontal plane. Two burr holes were drilled in the skull for bilateral injections of AAV5-EF1α-DIO-eNpHR3.0-mCherry (University of North Carolina Vector Core, 4×10^12 vg/ml) into the NAcSh (AP: +1.7; ML: ±0.9; DV: −7.2). Injectors were slowly lowered to the target depth and 0.5 µl of the viral vector was infused at a rate of 0.3 µl/min. Injectors were connected to 10 µl Hamilton syringes mounted on an infusion pump (Harvard Apparatus) via PE-20 tubing. After each injection, the injectors remained in place for an additional 5 minutes to allow diffusion of the virus in the brain tissue. Bone wax was placed in the burr holes and the skin incision was sutured closed. Rats were administered an additional 10 mL of saline and allowed to recover from surgery on a heating pad.

#### Behavioral experiment surgeries

For rats in behavioral experiments, an incision was made in the scalp and the skin was retracted with hemostats. After clearing the fascia, five to six small burr holes were made in the skull for placement of jeweler’s screws, with at least one positioned anterior to bregma and at least four positioned between bregma and lambda. Such a configuration was used to ensure that the cranial implant would be evenly secured across the skull. In Experiment 1 (i.e., optogenetic manipulation of BLA→NAcSh), the skull was leveled and then two additional burr holes were drilled in the skull for bilateral injections of AAV5-CAMKIIα-eNpHR3.0-mCherry (University of North Carolina Vector Core; 5.8×10^12 vg/ml) or AAV5-CAMKIIα-mCherry (4.9×10^12 vg/ml) in the BLA (AP: −3.3; ML: ±4.9; DV: −8.5, −8.1 from skull surface). Injectors were slowly lowered to the target depth and the viral vector was infused at a rate of 0.3 µl/min (0.4 µl at the most ventral DV coordinate and 0.2 µl at the more dorsal DV coordinate). Injectors were left in place for an additional 5 minutes after each injection to allow diffusion of the virus through the brain tissue. After viral injections were completed, bone wax was placed into each burr hole. Two more burr holes were then created for bilateral implantation of guide cannulae (22 gauge, Plastics One) above the NAcSh (AP: +1.5; ML: ±3.1; DV: −6.4 from skull surface at a 20° angle). Once both cannulae were lowered to their target depth, a thin layer of Metabond (Parkell) was applied over the skull and jeweler’s screws. After the Metabond had cured, dental cement was then used to anchor the guide cannulae in place. Sterile stylets were inserted into the guide cannulae and the incision around the cranial implant was sutured closed. Once Vetricyn antibacterial ointment was applied to the incision, rats were given an additional 10 mL of saline and allowed to recover from surgery on a heating pad. Rats were allowed to recover for 1 week before being food-restricted in preparation for behavioral training.

In Experiment 2 (i.e., optogenetic manipulation of D2R-expressing neurons in the NAcSh), the surgical procedures were identical to those described above, with the exception that viral infusions were in the NAcSh. After leveling the skull, two burr holes were drilled for bilateral implantation of guide cannulae above the NAcSh (AP: +1.5; ML: ±3.1; DV: −6.4 from skull surface at a 20° angle). Once the cannulae were secured with dental cement, injectors were inserted into each cannulae through which AAV5-EF1α-DIO-eNpHR3.0-mCherry (University of North Carolina Vector Core; 4×10^12 vg/ml) or AAV-EF1α-DIO-mCherry (5.3×10^12 vg/ml) was injected into the NAcSh (0.5 µl) at a rate of 0.3 µl/min. Injectors remained in place for an additional 5 minutes and were then replaced with sterile stylets. The incision around the cranial implant was sutured closed and Vetricyin was applied to the skin around the implant. Rats were administered 10 mL of saline and were placed on a heating pad to recover. After one week of recovery from surgery, rats were food-restricted in preparation for behavioral training.

### In vitro *electrophysiology*

#### Brain slice preparation

Rats were anesthetized using ketamine/xylazine cocktail (100 mg/kg ketamine, 10 mg/kg xylazine) to achieve a surgical plane of anesthesia (evaluated by the absence of response to tail or hind paw pinch) and transcardially perfused with ice-cold sucrose-laden artificial cerebrospinal fluid (ACSF) containing (in mM): 205 sucrose, 10 dextrose, 1 MgSO_4_, 2 KCl, 1.25 NaH_2_PO_4_, 1 CaCl_2_, and 25 NaHCO_3_, oxygenated with 95% O_2_/5% CO_2_. Rats were decapitated using a small animal guillotine, brains were extracted and submerged in perfusion solution, and then sectioned horizontally to produce 300 µm slices using a Leica VT1000 vibratome. Slices were transferred to incubation ACSF containing (in mM): 124 NaCl, 10 dextrose, 3 MgSO_4_, 2.5 KCl, 1.23 NaH_2_PO_4_, 1 CaCl_2_, and 25 NaHCO_3_, oxygenated with 95% O_2_/5% CO_2_ and maintained at 35 °C. After 30 minutes, the incubation chamber was permitted to passively equilibrate to room temperature for at least 30 minutes before recording. Recordings were performed in ACSF containing (in mM): 126 NaCl, 11 dextrose, 1.5 MgSO_4_, 3 KCl, 1.2 NaH_2_PO_4_, 2.4 CaCl_2_, and 25 NaHCO_3_, oxygenated with 95% O_2_/5% CO_2_, and maintained at 28 °C flowing at 2 mL/min through a perfusion chamber. Patch pipettes with an open tip resistance of 4-6 MΩ were prepared from borosilicate glass (Sutter Instrument BF150-86-10) using a Flaming/Brown pipette puller (Sutter Instrument SU-P97). Cells were visualized with an Olympus BX51WI upright stereomicroscope using IR-DIC optics, a 12-bit CCD camera (QImaging Rolera-XR), and µManager software (Edelstein et al., 2010). A TTL-controlled LED light source (X-Cite 110LED, Excelitas Technologies) paired with a green emission filter (Omega Optical XF414) was used to deliver green light to the sample through a 40x water-immersion objective lens. Electrophysiological recordings were performed using a CV7-B patch-clamp headstage, MultiClamp 700B amplifier, DigiData 1440A digital acquisition system, and pClamp 11 software (Axon Instruments/Molecular Devices). Data were acquired at 20 kHz. Voltage clamp recordings were lowpass filtered using a 2kHz Bessel filter. Patch pipettes were filled with a physiological internal solution containing (in mM): 125 potassium gluconate, 10 phosphocreatine, 1 MgCl_2_, 10 HEPES, 0.1 EGTA, 2 Na_2_ATP, 0.25 Na_3_GTP, adjusted to pH 7.25 and 295 mOsm. Data are presented uncorrected for the liquid junction potential.

#### Electrophysiology procedures

Neurons expressing eNpHR3.0 were targeted for whole-cell recording by their mCherry fluorescence. Immediately following establishment of whole-cell configuration, passive membrane properties (holding current, membrane resistance, and whole-cell capacitance) were evaluated in voltage clamp using a brief −10 mV hyperpolarizing step from a holding potential of −70 mV. Sensitivity to green light was evaluated in current-clamp configuration at the resting potential. Light-induced hyperpolarization was quantified as the mean voltage over a 500-millisecond period of green light exposure relative to the resting voltage, which was measured over a 1-second period before onset of the green light. To demonstrate eNpHR3.0-mediated suppression of firing, neurons were depolarized above action potential threshold using continuous positive current injection and 1-second green light was applied to demonstrate reduction or elimination of firing. In a subset of experiments, the D2R agonist quinpirole (Tocris #1061) was delivered into the perfusion system using a syringe pump to achieve a bath concentration of 10 µM. Changes in holding current required to voltage-clamp neurons at −70 mV were monitored and the effect of quinpirole was reported as the mean holding current observed during a 2-minute period during quinpirole exposure minus the mean holding current observed in a 3-minute baseline period immediately before application. Membrane resistance was evaluated from the same voltage-clamp step using the same time periods and is reported as the resistance during quinpirole exposure normalized to the baseline resistance. Effects of quinpirole on action potential firing frequency were evaluated using a current-clamp protocol that applied a 1-second excitatory current pulse (sufficient to evoke 5 or more action potentials in baseline conditions) every 10 seconds. Quinpirole-induced changes in firing frequency were evaluated in a 2-minute period during quinpirole exposure normalized to a baseline period. For all electrophysiological experiments, significance of effects observed in baseline subtracted or normalized data was evaluated using a one-sample Student’s *t*-test (with null hypothesis that mean = 0, or 1, respectively).

### Behavioral procedures

Before training in the Risky Decision-making Task (RDT), rats were shaped to perform separate components of the task, such as lever pressing and nosepoking to initiate trials as described previously (Orsini et al., 2017; Hernandez et al., 2019). Briefly, rats were first trained to associate the sound of a food pellet being deposited in the food trough with food delivery, after which they learned to press each of the two levers to receive a single food reward. Finally, rats were trained to nosepoke into the nosepoke hole to trigger the extension of a single lever, a press on which resulted in the delivery of one food pellet. Upon completing these shaping protocols, rats began training in a Reward Discrimination task. This task consisted of three blocks of 28 trials and lasted 56 minutes. The beginning of a trial was signaled with the illumination of the houselight and nosepoke light. A nosepoke into the nosepoke hole extinguished the nosepoke light and triggered the extension of one lever (forced choice trial; randomly presented) or both levers (free choice trial). If rats failed to nosepoke within 10 seconds, both lights were extinguished and the trial was scored as an omission. A lever press on one lever resulted in the delivery of 1 food pellet (small reward) whereas a press on the other lever resulted in the delivery of 2 food pellets (large reward). The identity of the lever (small vs. large reward) was counterbalanced across rats and sexes and remained consistent throughout the entire experiment. Failure to press levers within 10 seconds led to the retraction of the levers and the termination of lights in the chamber, and the trial was scored as an omission. Once a lever press occurred, levers were retracted and the light in the food trough was illuminated and remained so until food was collected by the rat or after 10 seconds had elapsed, whichever occurred first. Each block of 28 trials began with 8 forced choice trials in which a single lever was extended into the chamber (four presentations of each lever, randomized across the eight trials) and ended with 20 free choice trials in which both levers were extended into the chamber and rats were free to choose between them. Unlike the RDT (see below), each block was identical in structure and served to teach the rat about the overall design of the task before footshock punishment was introduced. Rats were trained on the Reward Discrimination task until they selected the large reward on at least 80% of the free choice trials for 2 consecutive days, after which they progressed to training in the RDT. On average, rats required no more than 5 sessions to reach these criteria.

The task structure for the RDT was identical to that of the Reward Discrimination task, with the exception that delivery of the large reward was accompanied by the possibility of a mild footshock punishment (Figure 1A). The probability of footshock delivery was 0% in the first block of trials and increased to 25% and 75% in the second and third block, respectively. Rats learned which probability of footshock delivery was in effect for each block during the forced choice trials that preceded the free choice trials. In the forced choice trials, the probability of receiving a footshock was dependent across the four trials in which the large, “risky” lever was available. For example, in the 25% trial block, one and only one of the forced choice trials would lead to the delivery of a footshock. In contrast, in the 75% trial block, three out of the four forced choice trials would result in the delivery of a footshock. In the free choice trials, however, the probability of footshock on a single trial was independent of the outcomes on other trials within that trial block. Hence, the probability of shock delivery was equivalent across all free choice trials, irrespective of whether previous trials resulted in shock deliveries. Shock intensities were initially set at 0.15 mA for females and 0.25 for males, but were individually adjusted for each rat during training to ensure that their baseline performance was in the middle of the parametric space prior to optogenetic manipulations. Shock intensities never exceeded 0.55 mA and never fell below 0.075 mA. Rats were trained on the RDT until behavioral stability was achieved (see *Experimental design and statistical analyses* for definition of behavioral stability).

**Figure 1.**
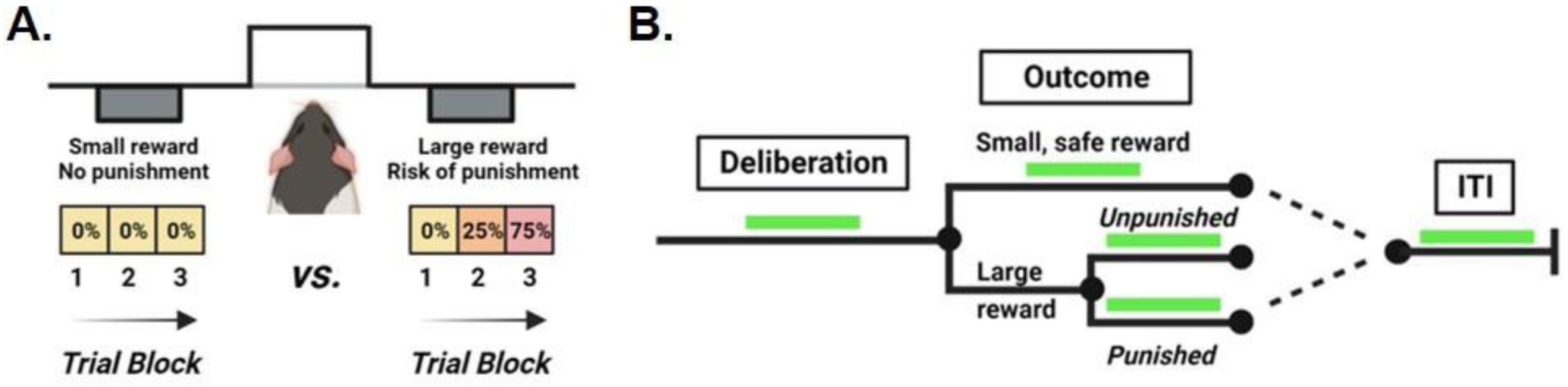
Schematic of The Risky Decision-making Task and timing of light delivery. **A.** In the Risky Decision-making Task (RDT), rats choose between two levers that differ in their reward magnitude and the risk of punishment. A press on one lever yields a small reward with no risk of punishment whereas a press on the other lever yields a large reward associated with risk of punishment. The probability of punishment delivery systematically increases across the 3 trial blocks. **B.** Light was delivered during discrete epochs of the decision process in separate test sessions. These epochs included 1) deliberation, 2) delivery of the small, safe reward, 3) delivery of the large, unpunished reward, 4) delivery of the large, punished reward, and 5) the intertrial interval (ITI). Using a within-subjects design, the order of the test sessions was counterbalanced across rats and successive optogenetic test sessions were separated by non-optogenetic test sessions to re-establish baseline performance.

Once behavioral stability on the RDT was reached, rats were lightly anesthetized and optic fibers were bilaterally inserted into the cannulae in the NAcSh, extending 1 mm beyond the tip of the cannulae. Fibers were cemented into place and dust caps were placed on the fibers to protect them from debris accumulation. From this point on, stainless steel patch cords were attached to the implanted fibers during each test session. Rats were tested in this manner to habituate them to being tethered prior to optogenetic manipulations. After behavioral stability was once again reached (2-3 days; tethering rats tended to induce variability in performance for several days), optogenetic manipulations began. In these sessions, laser delivery only occurred during free choice trials and was specific to distinct periods of the trial (Figure 1B): (1) deliberation (2) reward outcome, and (3) intertrial interval (ITI). On each free choice trial during deliberation sessions, the laser delivered light specifically during the period between the illumination of the nosepoke hole and a lever press. To ensure that full inhibition was in effect prior to the beginning of the deliberation phase, the onset of laser delivery preceded the illumination of the nosepoke hole by 0.5 seconds. If a rat did not press a lever within 10 seconds, light delivery was terminated. With respect to reward outcome optogenetic sessions, there were three different laser delivery sessions: (1) delivery of the small, safe reward (2) delivery of the large reward without punishment (large unpunished reward), and (3) delivery of the large reward with punishment (large punished reward). In each session, laser delivery was initiated as soon as a rat pressed a lever and was terminated 5 seconds after food delivery. Finally, serving as a behavioral control for laser delivery, the ITI optogenetic test session consisted of the delivery of laser light 8-15 seconds after each reward delivery, with each period of light delivery lasting 5 seconds. A randomized within-subjects design was used to determine the order in which the optogenetic test sessions were presented (across multiple days). After each optogenetic test session, rats were tethered and tested in the RDT until their behavior re-stabilized, after which they proceeded to the next optogenetic test session.

### Determination of shock intensity threshold

Upon completion of testing in the RDT, a subset of eNpHR3.0 rats in Experiment 1 (n=4) underwent a procedure to evaluate the shock intensities at which specific motor responses were generated. This assay, which was based on methods developed by Bonnet and Peterson (1975), was conducted twice for each rat over two consecutive days, with one session involving laser delivery (i.e., optogenetic inhibition) and the other involving no laser delivery (i.e., no optogenetic inhibition). Rats were placed in an operant chamber distinct from that in which they underwent RDT testing and given two minutes to acclimate to the chamber. A 0.40 mA unsignaled footshock (1 second) was then delivered to reduce spontaneous locomotor activity and facilitate observations of motor responses on subsequent trials at lower shock intensities. The shock intensity was then reduced to 0.05 mA and a series of five 1-second footshocks was delivered every 10 seconds. After each set of footshocks, the shock intensity was increased by 0.025 mA and another series of five footshocks were delivered. This pattern continued until all pre-determined motor responses of interest were identified. A shock intensity threshold for a specific motor response was established when three out of the five shock deliveries in a series elicited the specific motor response. The motor responses for which shock threshold intensities were determined were (1) flinch of one paw or a startle response (2) elevation of one or two paws (3) quick movement of more than two paws. During the session in which laser delivery occurred, light was delivered bilaterally through the implanted optic fibers with the same system and parameters (i.e., wavelength, light intensity) used for optogenetic manipulations during the RDT. Laser onset occurred concurrently with the onset of the footshock (1 second) and was terminated after an additional 4 seconds (5 seconds in total duration, which matches the duration of laser delivery during the large, punished reward session). During sessions in which no laser delivery occurred, rats were still tethered to equate behavioral conditions. The order of the test conditions (i.e., laser delivery vs. no laser delivery) was counterbalanced across rats.

### Fluorescence in situ hybridization (FISH), immunohistochemistry, and histology

#### FISH

To confirm that viral transduction was selective to neurons that expressed D2Rs within the NAcSh in D2-Cre transgenic rats, FISH (RNAscope, v1, ACDBio) combined with immunohistochemistry was performed on brain tissue from rats (n=4) that received viral infusions of AAV5-EF1α-DIO-eNpHR3.0-mCherry. Three weeks after surgery, rats were overdosed with intraperitoneal injections of Euthasol and transcardially perfused with ice-cold RNAase-free 0.9% NaCl followed by 4% paraformaldehyde (PFA) in 0.1M phosphate-buffered saline (PBS). Brains were extracted and post-fixed in 4% PFA for 12 hours, after which they were transferred to 30% sucrose in 0.1M PBS. Tissue was sectioned using a freezing cryostat at 12 µm and mounted directly onto Superfrost Plus Gold slides (Fisher Scientific) and then stored at −80°C. The modified procedure that combines RNAscope FISH with immunohistochemistry was previously described in Shallcross at el. (2019) with the following adjustments. The target probe used for detection of D2R mRNA was DRD2 (ACDBio; 315641-C2). Expression of mCherry was detected using anti-mCherry antibody (AbCam, ab167453, 1:1000) and visualized using donkey anti-rabbit secondary antibody conjugated to Alexa Fluor 488 (Life Technologies, A32790, 1:500). Finally, sections were counterstained with DAPI (Life Technologies), coverslipped using ProLong Gold antifade mounting reagent (Life Technologies) and sealed with nail polish.

D2R and mCherry signal (Z-stacks with 1µm step size) were acquired using Zeiss LSM70 confocal microscope and a 63X oil immersion objective. The procedures used for the quantification of D2R puncta and co-localization of D2R and mCherry signal has been described previously (Shallcross et al., 2019).

#### Immunohistochemistry

At the completion of behavioral testing, rats were overdosed with intraperitoneal injections of Euthasol and transcardially perfused with ice-cold 0.1M PBS, followed by 4% PFA in 0.1M PBS. Brain tissue was extracted, post-fixed in 4% PFA and then transferred to a 30% sucrose in 0.1M PBS solution. Tissue was sectioned (35 µm) on a cryostat maintained at −20°C. Sections were collected (1-in-6 series for Experiment 1; 1-in-4 series for Experiment 2) into wells filled with 0.1M PBS, after which they were transferred to cryoprotectant.

To confirm viral expression, immunohistochemistry was performed to amplify the mCherry fluorescent tag. Free-floating sections were washed three times for 10 minutes in 0.1M Tris-buffered saline (TBS) and then incubated in 0.1M TBS with 5% blocking buffer (Normal Donkey Serum, Jackson Immunoresearch, NC9624464) and 0.3% Triton-X 100 for 2 hours at room temperature. Tissue was then transferred into wells filled with primary antibody (rabbit anti-mCherry, AbCam, ab167453, 1:1000) diluted in 0.1M TBS with 5% blocking buffer and 0.3% Triton-X 100. Sections were incubated in the primary antibody solution for 72 hours at 4°C and then were washed three times for 10 minutes in 0.1M TBS. After the last wash, tissue was transferred into secondary antibody (donkey anti-rabbit conjugated to Alexa Fluor 488, Life Technologies, A32790, 1:300) diluted in 0.1M TBS with 5% blocking buffer and 0.3% Triton-X 100 and allowed to incubate in this solution for 2 hours at room temperature. Finally, tissue was washed three additional times for 10 minutes in 0.1M TBS and then mounted onto slides. Once sections on the slides were dry, slides were coverslipped with ProLong Gold Antifade mounting medium (Invitrogen, P36934) and sealed with nail polish. Processed tissue was visualized with a Zeiss AxioImager 2 microscope at 10X to confirm viral expression and determine optic fiber placement. Representative images were acquired using Zen Pro software at 10X with the same microscope.

### Experimental design and statistical analyses

Sample sizes for each group (eNpHR3.0 vs. mCherry) were determined *a priori* using power analyses in G*Power software. These analyses revealed that sample sizes of at least n=6 were necessary to detect significant differences between baseline and inhibition conditions in choice performance with effect sizes of ≥ 0.6 (assuming an α of 0.05). Additional rats were added to each group to account for potential attrition during the study and/or missed viral and/or cannula placements. Although both males and females were included in each experiment, group sizes were not powered to specifically test for sex differences as this was not the primary objective of the study. Sex was included as a between-subjects factor in all analyses, however, to inform experimental design for future studies (i.e., if there were a sex difference, future studies would be sufficiently powered to explicitly examine sex differences). In all analyses, a *p-value* ≤ 0.05 was considered statistically significant. If parent ANOVAs yielded main effects or significant interactions, additional *post-hoc* ANOVAs or *t*-tests were conducted to determine the source of the significance and *p-values* were adjusted to account for multiple comparisons (Bonferroni’s correction). Effect sizes are reported as ƞ^2^ for ANOVAs and as the absolute value of Cohen’s d for independent sample’s *t*-tests.

Raw files were extracted from Graphic State 4.0 using customized analysis templates and exported to Microsoft Excel. Excel was used to isolate behavioral measures of interest, including lever presses, latencies to lever press, latencies to initiate a trial (nosepoke), latencies to collect food rewards and omissions. Data were analyzed using SPSS 27.0 and figures were created using GraphPad 9.4.1. The primary dependent variable was the percentage of free choice trials on which a rat chose the large, risky reward. Individual rats were trained in the RDT until they displayed stable choice performance, which was defined as choice behavior with a coefficient of variation of ≤ 30% for percent choice of the large, risky reward in each block for a sliding window of at least two consecutive test sessions. Upon reaching stability, rats began the series of optogenetic test sessions. Following each optogenetic test session, rats were re-tested in the RDT until the stability criterion was once again achieved, after which they advanced to the next optogenetic test session. A two-factor repeated-measures ANOVA was used to assess the effects of optogenetic inhibition on choice behavior, with session (baseline versus inhibition) and trial block as within-subjects factors. For this and all subsequent analyses, sex was included as a between-subjects factor; if there was no main effect of sex or sex X session interaction, data were collapsed across sexes and analyzed and presented accordingly. If there was a main effect of session and/or a session X trial block interaction, trial-by-trial analyses were conducted to determine whether optogenetic inhibition altered the extent to which the outcome of a previous trial affected subsequent choice. Providing a measure of sensitivity to rewarding outcomes, win-stay trials were defined as trials on which a rat continued to choose the large, risky lever after receiving a large, unpunished reward. This variable was calculated by the dividing the number of trials on which the rat selected the large, risky lever after receipt of the large, unpunished reward by the total number of free choice trials on which the rat received the large, unpunished reward. In contrast, lose-shift trials provided a measure of sensitivity to negative feedback (i.e., punishment) and were defined as trials on which a rat shifted its choice to the small, safe lever after receiving a large, punished reward. This variable was calculated by dividing the number of trials on which the rat selected the small, safe lever after receipt of the large punished reward by the total number of free choice trials on which the rat received the large punished reward. Effects of optogenetic inhibition on win-stay and lose-shift trials were determined using a repeated-measures ANOVA, with trial type (win-stay versus lose-shift) and session (baseline versus inhibition) included as within-subjects factors.

If optogenetic manipulations altered choice behavior, ancillary behavioral measures, including latencies to lever press, nosepoke, and collect food rewards, were also compared between baseline and optogenetic test sessions. Latency to lever press during free choice trials was defined as the period of time between the nosepoke to trigger lever extension and a lever press. Because rats displayed a near-exclusive preference for the large reward in Block 1, latencies to press levers during free choice trials were constrained to Blocks 2 and 3 (i.e., there were insufficient data for latencies to press the small reward in Block 1, precluding its comparison to latencies to press the large reward). A repeated-measures ANOVA was used to assess the effects of optogenetic inhibition on latencies to press levers, with session (baseline versus inhibition), lever identity (small, safe lever versus large, risky lever) and trial block as within-subjects factors. Because onset of the laser began 0.5 seconds prior to the illumination of the nosepoke hole in test sessions in which the light was delivered during deliberation, latencies to nosepoke (i.e., the interval of time between the illumination of the nosepoke and a nosepoke) were also analyzed to determine whether light delivery altered this behavioral response. Specifically, a repeated-measures ANOVA was used to compare latencies to nosepoke between baseline and the deliberation test condition, with session and trial block as within-subjects factors. In sessions in which light was delivered during the reward delivery period, latencies to collect food were analyzed using a repeated-measures ANOVA, with session and trial block as within-subjects factors. Finally, omissions during free choice trials were calculated by dividing the number of omitted free choice trials by the total number of possible free choice trials (60) and multiplying this number by 100. For the large, punished reward optogenetic test session, analysis of omissions was limited to Blocks 2 and 3 as inhibition never occurred during Block 1. Omissions were then compared between baseline and optogenetic test sessions using a repeated-measures ANOVA (the inclusion of sex as a between-subjects factor necessitated the use of an ANOVA over a *t*-test). Notably, the percentage of omitted free choice trials was analyzed for each optogenetic manipulation, irrespective of whether there was an effect on choice behavior.

Shock intensity thresholds in eNpHR3.0 rats were compared between laser delivery and no laser delivery sessions using a repeated-measures ANOVA, with session and the type of motor response (one paw flinch, two paw flinches, rapid movement of more than two paws) included as within-subjects factors.

## Results

For behavioral experiments, descriptions of statistical results will be restricted to those directly related to the effects of optogenetic inhibition (or laser delivery in control rats). Unless otherwise noted, there were no sex differences in risk taking (collapsed across optogenetic inhibition/light delivery conditions; all *p-values* > 0.05). Similarly, for all analyses in which trial block was included as a within-subjects factor, there was a main effect of trial block (*p-values* < 0.05); this effect will therefore not be reported further.

### Experiment 1

#### Histology

A total of 54 Long-Evans rats (n=29, male; n=25, female) were used in Experiment 1. Of the 54, 29 rats (n=16, male; n=13, female) received intra-BLA infusions of the viral vector containing eNpHR3.0. Four females and four males in this group lost their cranial implants prior to any optogenetic manipulations and were therefore excluded from the study. In addition, three females and two males were excluded due to lack of viral transduction (i.e., no evidence of viral injection in the BLA and no terminal expression in the NAcSh). Twenty-five rats (n=13, male; n=12, female) received intra-BLA infusions of the viral vector containing mCherry. Five females and three males in this group lost their cranial implant prior to any optogenetic manipulations and were therefore excluded from the study. Three females and three males were also excluded as a result of poor viral transduction in the BLA and BLA terminals in the NAcSh. Consequently, the final sample sizes were n=16 for the eNpHR3.0 group (n=10, male; n=6, female) and n=11 for the mCherry group (n=7, male; n=4, female). Figure 2 depicts the placements of the optic fibers implanted in the NAcSh for rats in the eNpHR3.0 (Figure 2A) and control (Figure 2B) groups. Importantly, some rats did not undergo every optogenetic test condition due to illness or loss of their cranial implant.

**Figure 2.**
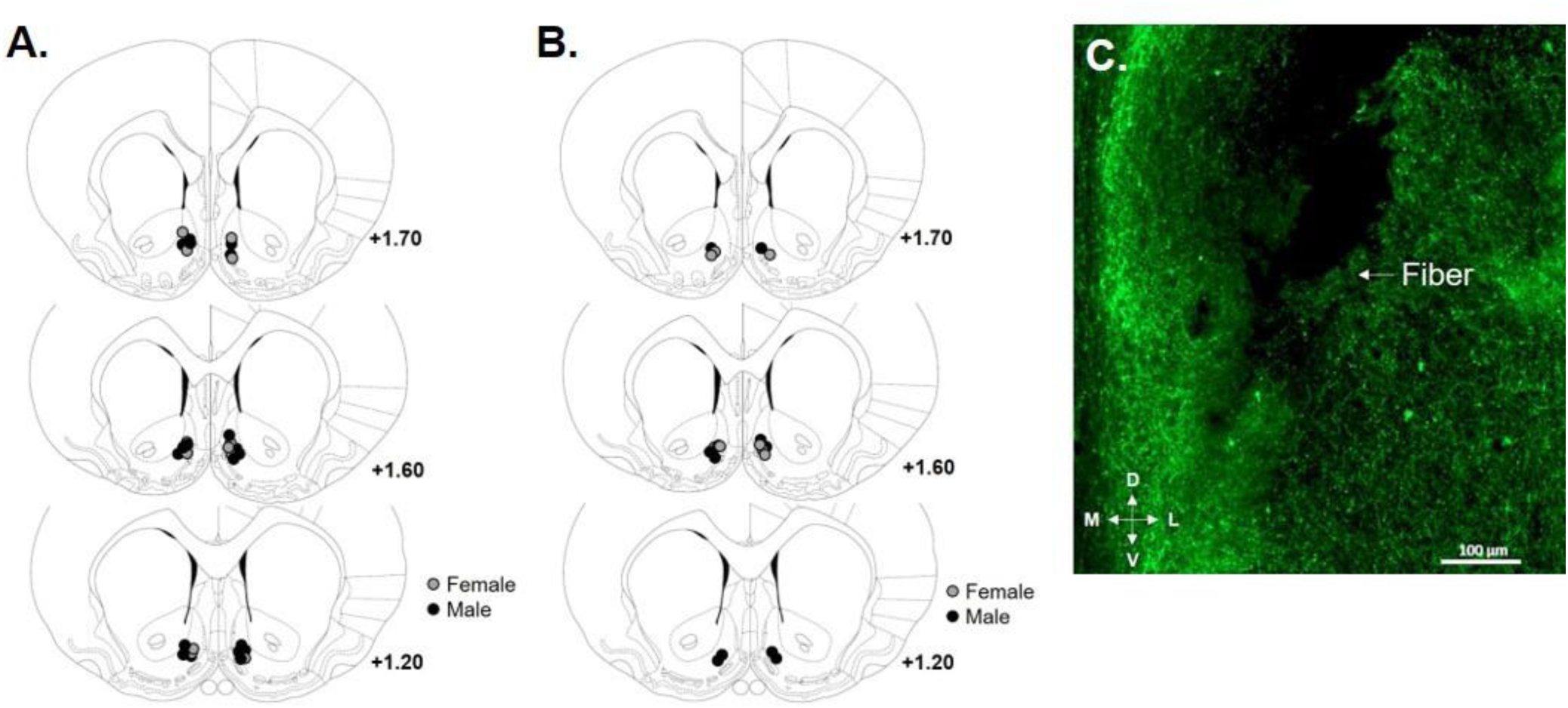
Optic fiber placement and BLA terminal expression of eNpHR3.0 in the NAcSh. **A.** Circles represent the location of the tip of the optic fiber in the nucleus accumbens shell (NAcSh) of rats with BLA terminal expression of eNpHR3.0. **B.** Circles represent the location of the tip of the optic fiber in the NAcSh of rats with mCherry (control group) terminal expression. **C.** Representative image of the tip of the optic fiber and BLA terminal expression of mCherry in the NAcSh of an eNpHR3.0 rat.

### Optogenetic inhibition of BLA terminals in the NAcSh during decision making in eNpHR3.0 rats

#### Inhibition during deliberation

Optogenetic inhibition of the BLA→NAcSh during deliberation (n=9, male, n=4, female) significantly increased choice of the large, risky reward (increased risk taking) in both males and females [Figure 3A; session, *F* (1, 11) = 5.80, *p* = 0.04, ƞ^2^ = 0.35; session X sex, *F* (1, 11) < 0.01, *p* = 0.98, ƞ^2^ < 0.01; session X trial block, *F* (2, 22) = 4.44, *p* = 0.02, ƞ^2^ = 0.29; session X sex X trial block, *F* (2, 22) < 0.01, *p* = 0.99, ƞ^2^ < 0.01]. Despite the effects of inhibition on choice performance, there was no effect of inhibition on the percentage of win-stay and lose-shift trials [Figure 3B; session, *F* (1, 11) = 0.41, *p* = 0.54, ƞ^2^ = 0.04; session X sex, *F* (1, 11) = 0.11, *p* = 0.74, ƞ^2^ = 0.01; session X trial type, *F* (1, 11) = 0.70, *p* = 0.42, ƞ^2^ = 0.06; session X sex X trial type, *F* (1, 11) = 0.03, *p* = 0.86, ƞ^2^ < 0.01]. To determine whether inhibition during deliberation had any lasting effect on task performance (beyond the session in which inhibition took place), baseline performance was compared to performance in the RDT 24 hours after the optogenetic test session using a three-factor repeated-measures ANOVA. This analysis revealed that there was no difference in performance between baseline and post-inhibition sessions [session, *F* (1, 10) = 0.88, *p* = 0.37, ƞ^2^ = 0.08; sex X session, *F* (1, 10) = 0.56, *p* = 0.47, ƞ^2^ = 0.05; session X trial block, *F* (2, 20) = 0.93, *p* = 0.41, ƞ^2^ = 0.09; session X sex X trial block, *F* (2, 20) = 1.02, *p* = 0.38, ƞ^2^ = 0.09]. Considered together, these results reveal that activity in the BLA→NAcSh during the deliberative part of the decision process biases subsequent choice toward small, safer options.

**Figure 3.**
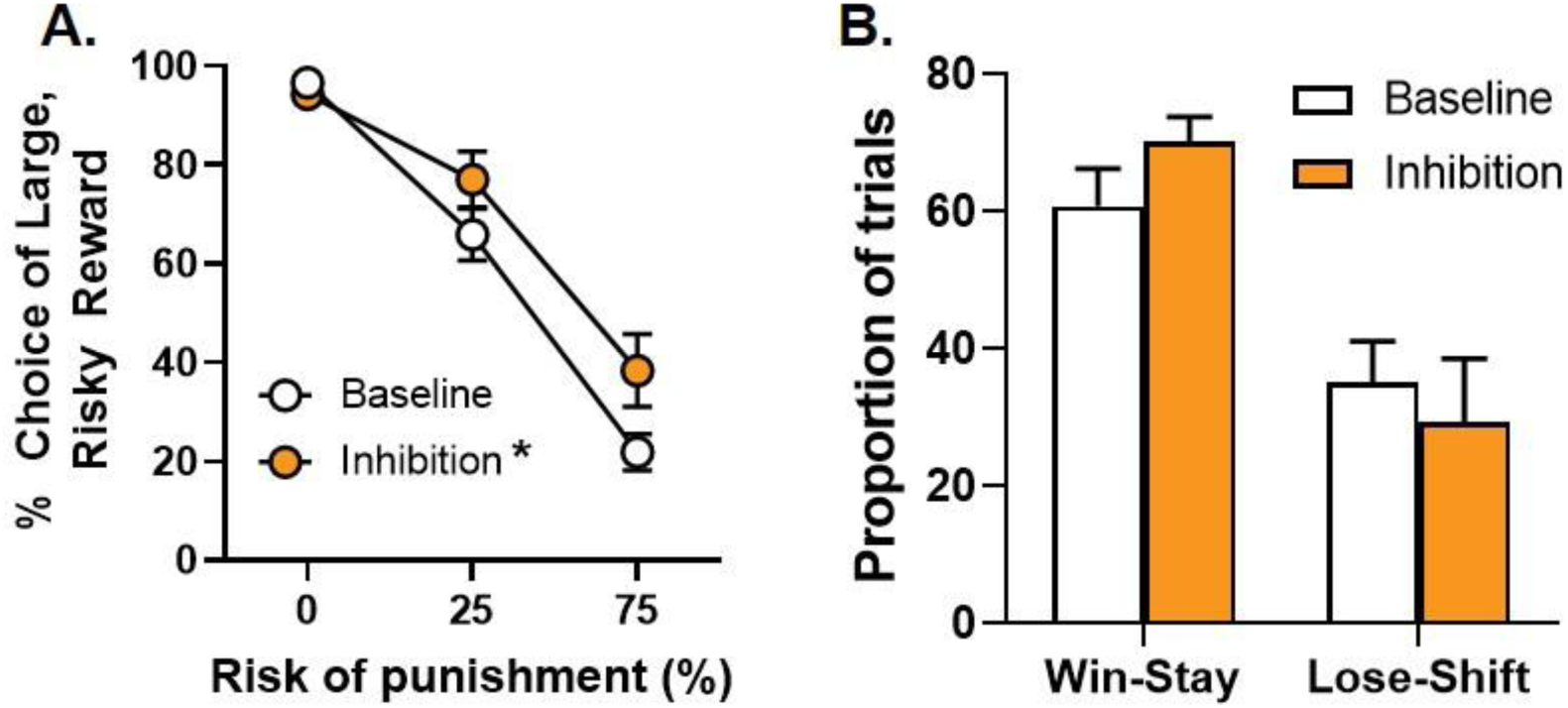
Optogenetic inhibition of BLA projections to the NAcSh during deliberation. **A.** Inhibition of the BLA→NAcSh during deliberation increased choice of the large, risky reward. **B.** There was no effect of inhibition of the BLA→NAcSh during deliberation on the percentage of win-stay or lose-shift trials. Data are represented as mean ± SEM. Asterisks indicate a significant difference between baseline (no laser delivery) and optogenetic inhibition (*p* < 0.05).

The onset of laser delivery during the deliberation test condition was 0.5 seconds prior to the start of each trial. Hence, to determine whether inhibition altered the latency to nosepoke to trigger lever insertion into the chamber, nosepoke latencies during free choice trials were compared between baseline sessions and the optogenetic test sessions. Despite an overall main effect of sex, with females taking longer to nosepoke than males [*F* (1, 11) = 7.35, *p* = 0.02, ƞ^2^ = 0.40], there was no effect of inhibition on this behavioral measure in males or females [session, *F* (1, 11) = 0.08, *p* = 0.79, ƞ^2^ < 0.01; session X sex, *F* (1, 11) = 1.29, *p* = 0.28, ƞ^2^ = 0.11; session X trial block, *F* (2, 22) = 1.13, *p* = 0.34, ƞ^2^ = 0.09; session X sex X trial block, *F* (2, 22) = 1.19, *p* = 0.32, ƞ^2^ = 0.10]. There was no effect of inhibition on latencies to press levers during the free choice trials [session, *F* (1, 8) = 0.10, *p* = 0.76, ƞ^2^ = 0.01; session X sex, *F* (1, 8) = 0.23, *p* = 0.64, ƞ^2^ = 0.03; session X lever identity (small, safe versus large, risky lever), *F* (1, 8) = 0.89, *p* = 0.37, ƞ^2^ = 0.10; session X lever identity X trial block, *F* (1, 8) = 1.11, *p* = 0.32, ƞ^2^ = 0.12; session X sex X trial block, *F* (1, 8) = 1.59, *p* = 0.24, ƞ^2^ = 0.17; session X sex X lever identity X trial block, *F* (1, 8) = 1.99, *p* = 0.20, ƞ^2^ = 0.20]. Although this analysis yielded a significant interaction between session, sex and lever identity [*F* (1, 8) = 5.81, *p* = 0.04], the results of *post-hoc* ANOVAs comparing latencies to press each lever between sessions separately for males and females did not reach statistical significance when correcting for multiple comparisons. Finally, optogenetic inhibition of the BLA→NAcSh did not impact the percentage of omitted free choice trials [Table 1; session, *F* (1, 11) < 0.01, *p* = 0.96, ƞ^2^ < 0.01; session X sex, *F* (1, 11) = 1.73, *p* = 0.22, ƞ^2^ = 0.14].

**Table 1.** Mean (± standard error of the mean) omissions.

|  | *eNpHR3.0* |  | *mCherry* |  |
| --- | --- | --- | --- | --- |
|  | Male | Female | Male | Female |
| **Experiment 1** |  |  |  |  |
| *Deliberation* |  |  |  |  |
| *Baseline* | 0.66 (0.46) | 2.43 (1.20) | 1.66 (0.70) | 5.83 (1.14) |
| *Inhibition* | 1.94 (1.00) | 1.94 (1.00) | 3.49 (1.54) | 5.21 (2.60) |
| *Delivery of small, safe reward* |  |  |  |  |
| *Baseline* | 0.93 (0.63) | 3.53 (1.58) | N/A | N/A |
| *Inhibition* | 1.51 (0.54) | 5.83 (3.94) | N/A | N/A |
| *Delivery of large, unpunished reward* |  |  |  |  |
| *Baseline* | 0.83 (0.24) | 5.07 (1.96) | N/A | N/A |
| *Inhibition* | 0.25 (0.25) | 7.17 (2.52) | N/A | N/A |
| *Delivery of large, punished reward* |  |  |  |  |
| *Baseline* | 3.02 (2.16) | 9.19 (3.67) | 1.69 (0.89) | N/A |
| *Inhibition* | 13.33 (12.39) | 3.44 (0.59) | 3.19 (1.81) | N/A |
| *Intertrial interval* |  |  |  |  |
| *Baseline* | 0.76 (0.23) | 6.93 (2.85) | N/A | N/A |
| *Inhibition* | 1.57 (0.69) | 6.80 (3.47) | N/A | N/A |
| **Experiment 2** |  |  |  |  |
| *Deliberation* |  |  |  |  |
| *Baseline* | 1.47 (0.60) | 12.44 (4.53) | 2.87 (1.30) | 4.92 (1.36) |
| *Inhibition* | 2.78 (2.44) | 10.14 (6.09) | 3.09 (1.60) | 6.99 (1.89) |
| *Delivery of small, safe reward* |  |  |  |  |
| *Baseline* | 3.80 (1.82) | 7.67 (3.90) | 6.25 (1.25) | 4.01 (1.40) |
| *Inhibition* | 4.31 (2.50) | 7.17 (2.78) | 2.50 (2.50) | 7.08 (3.48) |
| *Delivery of large, unpunished reward* |  |  |  |  |
| *Baseline* | 1.29 (1.23) | 10.64 (5.21) | N/A | N/A |
| *Inhibition* | 1.33 (0.57) | 10.83 (2.50) | N/A | N/A |
| *Delivery of large, punished reward* |  |  |  |  |
| *Baseline* | 0.69 (0.33) | 6.36 (3.89) | N/A | N/A |
| *Inhibition* | 0.00 (0.00) | 13.50 (6.75) | N/A | N/A |
| *Intertrial interval* |  |  |  |  |
| *Baseline* | 1.62 (1.53) | 8.87 (4.61) | N/A | N/A |
| *Inhibition* | 1.67 (0.60) | 9.03 (2.73) | N/A | N/A |

#### Inhibition during delivery of the small, safe reward

A two-factor, repeated-measures ANOVA revealed that optogenetic inhibition of the BLA→NAcSh during delivery of the small, safe reward (n=9, male; n=5, female) had no effect on choice of the large, risky reward [Figure 4A; session, *F* (1, 12) = 1.64, *p* = 0.23, ƞ^2^ = 0.12; session X sex, *F* (1, 12) = 0.86, *p* = 0.37, ƞ^2^ = 0.07; session X trial block, *F* (2, 24) = 2.40, *p* = 0.11, ƞ^2^ = 0.17; session X sex X trial block, *F* (2, 24) = 0.80, *p* = 0.46, ƞ^2^ = 0.06]. Consistent with a lack of an effect on choice behavior, there was no effect of inhibition on percentage of omitted free choice trials [Table 1; session, *F* (1, 12) = 1.28, *p* = 0.28, ƞ^2^ = 0.10; session X sex, *F* (1, 12) = 0.45, *p* = 0.56, ƞ^2^ = 0.04]. Collectively, these data reveal that activity in the BLA→NAcSh is not necessary for evaluation of small, safe rewards to guide subsequent risk-taking behavior.

**Figure 4.**
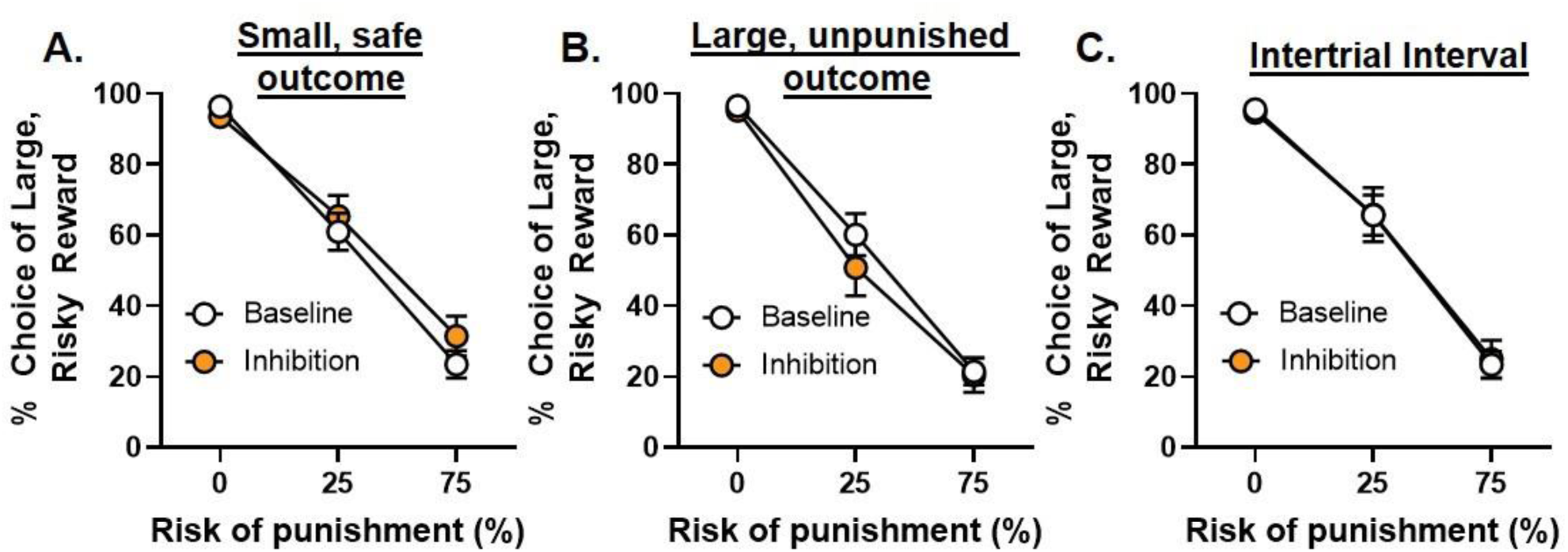
Optogenetic inhibition of BLA projections to the NAcSh during other phases of the RDT. **A.** There was no effect of BLA→NAcSh inhibition during delivery of the small, safe reward on choice of the large, risky reward. **B.** There was no effect of BLA→NAcSh inhibition during delivery of the large, unpunished reward on choice of the large, risky reward. **C.** There was no effect of BLA→NAcSh inhibition during the intertrial interval on choice of the large, risky reward. Data are represented as mean ± SEM.

#### Inhibition during delivery of the large, unpunished reward

Optogenetic inhibition of the BLA→NAcSh during delivery of the large, unpunished reward (n=9, male; n=5, female) did not affect choice of the large, risky reward [Figure 4B; session, *F* (1, 12) = 2.15, *p* = 0.17, ƞ^2^ = 0.15; session X trial block, *F* (2, 24) = 2.31, *p* = 0.12, ƞ^2^ = 0.16]. Despite a main effect of sex (i.e., greater risk taking in males; *F* (1, 12) = 15.29, *p* < 0.01, ƞ^2^ = 0.56], there were no significant session X sex [*F* (1,12) = 1.10, *p* = 0.32, ƞ^2^ = 0.08] or session X sex X trial block [*F* (2, 24) = 1.46, *p* = 0.25, ƞ^2^ = 0.11] interactions, indicating that inhibition was ineffective in altering risk taking in males or females. Although females omitted significantly more free trials overall [*F* (1, 12) = 15.98, *p* < 0.01, ƞ^2^ = 0.57], there was no main effect of session [*F* (1, 12) = 0.71, *p* = 0.42, ƞ^2^ = 0.06] nor a significant sex X session interaction [*F* (1, 12) = 2.22, *p* = 0.16, ƞ^2^ = 0.16] on percentage of omitted free choice trials (Table 1). Together, these results indicate that activity in the BLA→NAcSh is not necessary for evaluation of large, unpunished rewards to guide subsequent risk-taking behavior.

### Inhibition during delivery of the large, punished reward

In contrast to the effects of inhibition during the large, unpunished reward, optogenetic inhibition of the BLA→NAcSh during the delivery of the large, punished reward (n=8, male; n=4, female) increased choice of the large, risky reward [session, *F* (1, 10) = 5.70, *p* = 0.04, ƞ^2^ = 0.36; session X trial block, *F* (2, 20) = 11.19, *p* < 0.01, ƞ^2^ = 0.53]. Unexpectedly, in addition to a main effect of sex [*F* (1, 10) = 7.53, *p* = 0.02, ƞ^2^ = 0.43], there were significant session X sex [*F* (1, 10) = 13.96, *p* < 0.01, ƞ^2^ = 0.58] and session X sex X trial block [*F* (2, 20) = 9.67, *p* < 0.01, ƞ^2^ = 0.49] interactions. To determine the source of the significant interaction, subsequent two-factor repeated-measures ANOVAs were conducted to compare performance between baseline and inhibition sessions separately for males and females. These analyses revealed that optogenetic inhibition increased risk taking in males [Figure 5A; session, *F* (1, 7) = 31.85, *p* = 0.01, ƞ^2^ = 0.82; session X trial block, *F* (2, 14) = 40.09, *p* < 0.01, ƞ^2^ = 0.85], but did not affect performance in females [Figure 5B; session, *F* (1, 3) = 0.56, *p* = 0.52, ƞ^2^ = 0.15; session X trial block, *F* (2, 6) = 1.89, *p* = 0.23; ƞ^2^ = 0.39]. Although inhibition increased risk taking in males, there was no effect of this manipulation on the percentage of win-stay and lose-shift trials [Figure 5C; session, *F* (1, 7) = 3.05, *p* = 0.12, ƞ^2^ = 0.30; session X trial type, *F* (1, 7) = 2.09, *p* = 0.19, ƞ^2^ = 0.23]. Effects of inhibition on risk taking in males did not persist beyond the optogenetic session as there were no differences between baseline performance and performance on the RDT 24 hours after the manipulation [session, *F* (1, 7) = 0.95, *p* = 0.37, ƞ^2^ = 0.12; session X trial block, *F* (2, 14) = 1.15, *p* = 0.35, ƞ^2^ = 0.14]. Collectively, these data show that activity of the BLA→NAcSh in males, but not females, is necessary for evaluation of large rewards that are accompanied by punishment to guide subsequent choice toward safer options.

**Figure 5.**
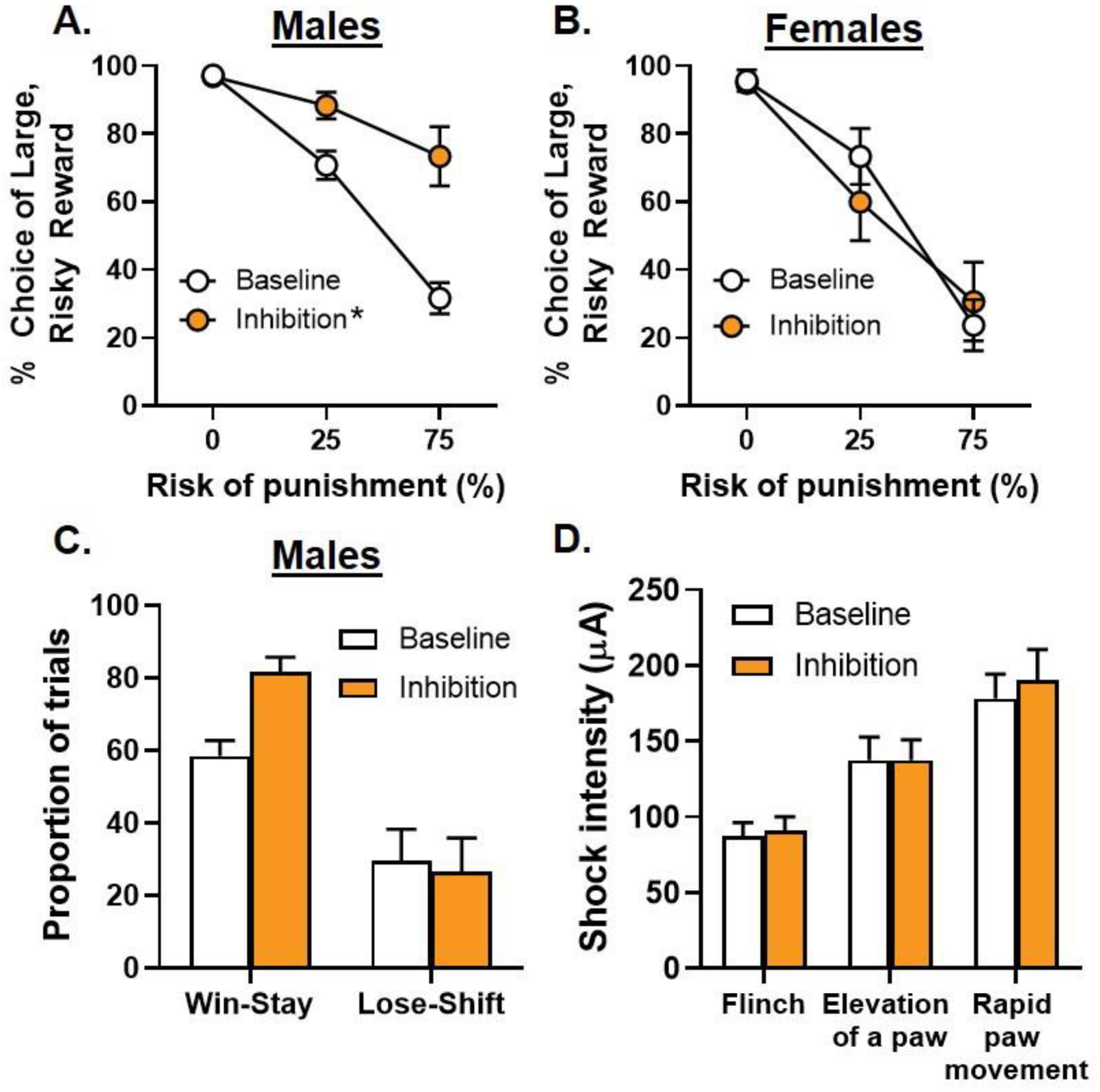
Optogenetic inhibition of BLA projections to the NAcSh during delivery of the large, punished reward. **A.** Inhibition of the BLA→NAcSh during delivery of the large, punished reward increased choice of the large, risky reward in males. **B.** There was no effect of BLA→NAcSh inhibition on choice of the large, risky reward in females. **C.** There was no effect of BLA→NAcSh inhibition on the percentage of win-stay and lose-shift trials in males. **D.** Inhibition of the BLA→NAcSh did not alter shock intensity thresholds at which shock delivery elicited a paw flinch, elevation of 1 or 2 paws or rapid movement of all 4 paws. Data are represented as mean ± SEM. Asterisks indicate a significant difference between baseline (no laser delivery) and optogenetic inhibition (*p* < 0.05).

To determine whether inhibition affected other behavioral measures in males, latencies to press levers during free choice trials and collect food were analyzed. These analyses were constrained to data from the blocks in which inhibition occurred (i.e., Blocks 2 and 3). Analysis of latencies to press levers revealed no effect of inhibition on latencies to press either lever [session, *F* (1, 6) = 0.65, *p* = 0.45, ƞ^2^ = 0.10; session X lever identity, *F* (1, 6) = 3.59, *p* = 0.11, ƞ^2^ = 0.37; session X lever identity X trial block, *F* (1, 6) = 0.11, *p* = 0.75, ƞ^2^ = 0.02]. Similarly, latencies to collect food were also not impacted by optogenetic inhibition [session, *F* (1, 6) = 0.79, *p* = 0.41, ƞ^2^ = 0.12; session X trial block, *F* (1, 6) = 0.08, *p* = 0.79, ƞ^2^ = 0.01]. Finally, inhibition during delivery of the large, risky reward had no effect on the percentage of omitted free trials in Blocks 2 and 3 in males [Table 1; *t* (7) = −0.81, *p* = 0.45, d = 0.29] or in females [*t* (3) = 1.38, *p* = 0.26, d = 0.69].

#### Inhibition during the intertrial interval

To confirm that the effects of inhibition of the BLA→NAcSh during deliberation and delivery of the large, punished reward were not due to non-specific effects of eNpHR3.0 activation, light was delivered during the intertrial interval (ITI; n=9, male; n=4, female). Unsurprisingly, there were no effects of optogenetic inhibition on choice of the large, risky reward [Figure 4C; session, *F* (1, 11) = 0.11, *p* = 0.75, ƞ^2^ = 0.01; session X sex, *F* (1, 11) = 1.38, *p* = 0.27, ƞ^2^ = 0.11; session X trial block, *F* (2, 22) = 0.02, *p* = 0.98, ƞ^2^ < 0.01; session X sex X trial block, *F* (2, 22) = 0.34, *p* = 0.72, ƞ^2^ = 0.03]. Similarly, although females made more omissions overall [Table 1; *F* (1, 11) = 8.33, *p* = 0.02, ƞ^2^ = 0.43], there was no effect of inhibition during the ITI on the percentage of omitted free choice trials [session, *F* (1, 11) = 0.17, *p* = 0.69, ƞ^2^ = 0.02; session X sex, *F* (1, 11) = 0.31, *p* = 0.59, ƞ^2^ = 0.03].

#### Inhibition during assessment of shock intensity thresholds

One interpretation of the effects of inhibition of the BLA→NAcSh during the delivery of the large, punished reward is that inhibition decreased sensitivity to footshock, and that the increase in risk taking was therefore secondary to this effect. To address this possibility, a subset of male rats (n=4) was tested in a behavioral assay used to determine thresholds at which specific motor responses were elicited by the delivery of footshock (see *Materials and Methods* for a description of the assay). Because the effects of inhibition on risk taking were only observed in males, female rats were not included in this part of the experiment. Results of a three-factor repeated-measures ANOVA showed that shock intensity thresholds for each of the pre-selected motor responses did not differ between laser delivery and no laser delivery sessions [Figure 5D; session, *F* (1, 3) = 2.78, *p* = 0.19, ƞ^2^ = 0.48; session X motor response, *F* (2, 6) = 0.62, *p* = 0.57, ƞ^2^ = 0.17]. Hence, effects of inhibition during delivery of the large, punished reward cannot be attributed to inhibition-induced alterations in sensitivity to footshock.

### Light delivery to BLA terminals in the NAcSh during decision making in control rats

To confirm that light delivery alone did not cause the observed changes in risk taking (e.g., via local tissue heating), another group of rats received intra-BLA injections of an AAV containing mCherry alone and were implanted with optic fibers in the NAcSh (n= 7, male; n= 4, female). They were then trained in the RDT and, upon achieving stable baseline performance, underwent optogenetic test sessions. These test sessions were restricted to those in which optogenetic inhibition of the BLA→NAcSh in the eNpHR3.0 group significantly altered risk taking (i.e., deliberation and delivery of the large, punished reward). The order of optogenetic test sessions was counterbalanced across rats, and these sessions were separated by non-optogenetic test sessions to re-establish baseline performance.

#### Inhibition during deliberation

Light delivery during deliberation (n=7, male; n=4, female) did not affect choice of the large, risky reward [Figure 6A; session, *F* (1, 9) = 2.55, *p* = 0.15, ƞ^2^ = 0.22; session X sex, *F* (1, 9) = 0.1.25, *p* = 0.29, ƞ^2^ = 0.12; session X trial block, *F* (2, 18) = 1.26, *p* = 0.31, ƞ^2^ = 0.12; session X sex X trial block, *F* (2, 18) = 0.95, *p* = 0.41, ƞ^2^ = 0.10]. There was also no effect of light delivery on latency to nosepoke to trigger lever extension [session, *F* (1, 9) = 2.43, *p* = 0.15, ƞ^2^ = 0.21; session X sex, *F* (1, 9) = 1.64, *p* = 0.23, ƞ^2^ = 0.15; session X trial block, *F* (2, 18) = 0.05, *p* = 0.95, ƞ^2^ < 0.01; session X sex X trial block, *F* (2, 18) = 2.46, *p* = 0.11, ƞ^2^ = 0.22]. Similarly, light delivery did not affect latencies to press levers during free choice trials [session, *F* (1, 8) = 0.08, *p* = 0.79, ƞ^2^ = 0.01; session X sex, *F* (1, 8) = 1.18, *p* = 0.31, ƞ^2^ = 0.13; session X lever identity, *F* (1, 8) = 3.09, *p* = 0.12, ƞ^2^ = 0.28; session X lever identity X sex, *F* (1, 8) = 1.96, *p* = 0.20, ƞ^2^ = 0.20; session X lever identity X trial block, *F* (1, 8) = 1.90, *p* = 0.21, ƞ^2^ = 0.19; session X lever identity X sex X trial block, *F* (1, 8) = 2.44, *p* = 0.16, ƞ^2^ = 0.23]. Lastly, light delivery during deliberation did not affect the percentage of omitted free choice trials [Table 1; session, *F* (1, 9) = 0.18, *p* = 0.69, ƞ^2^ = 0.02; session X sex, *F* (1, 9) = 0.73, *p* = 0.41, ƞ^2^ = 0.08]. Considered together, these results indicate that light delivery alone did not account for the increased risk taking that was observed when the BLA→NAcSh was inhibited during deliberation.

**Figure 6.**
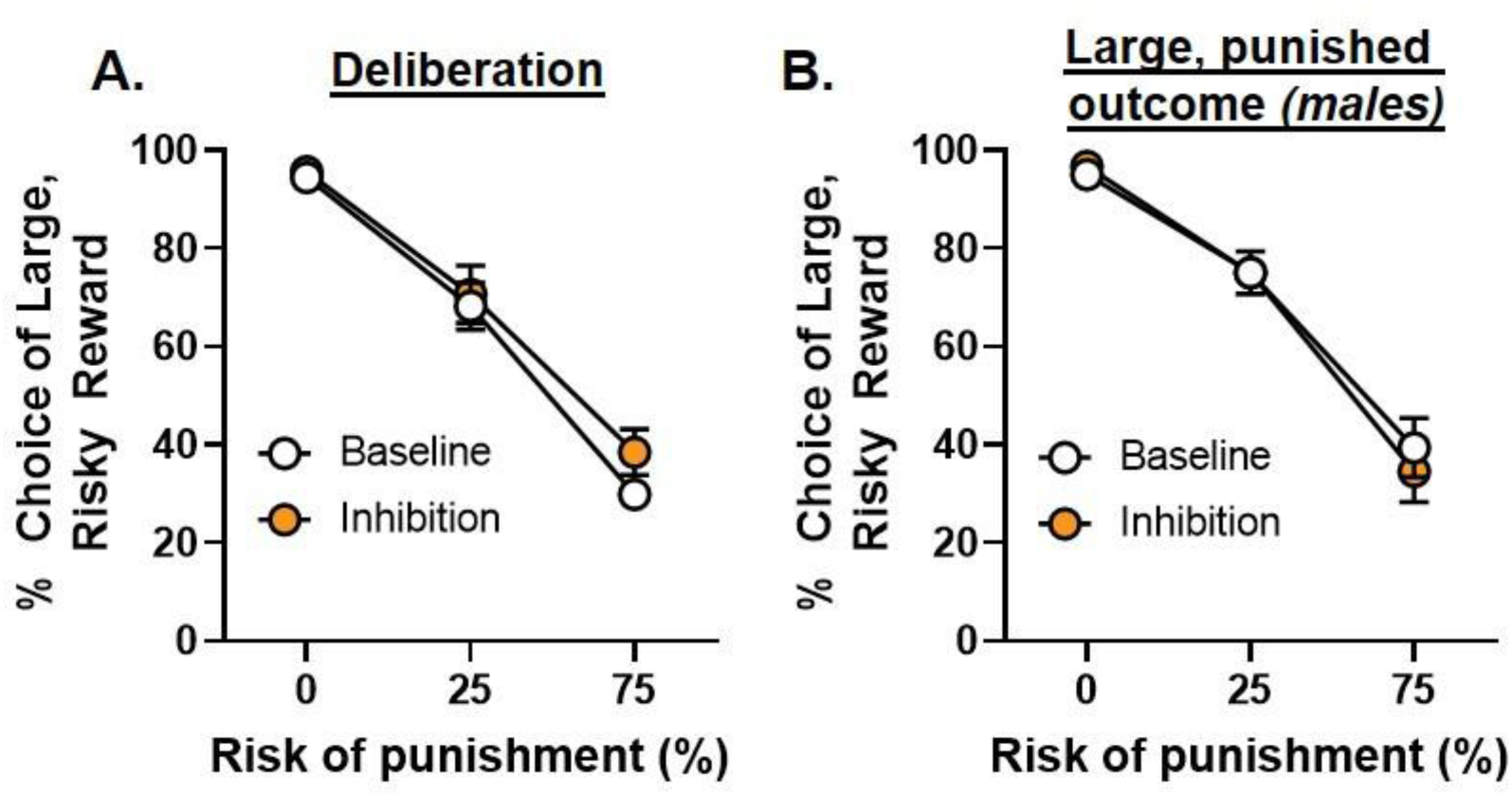
Light delivery to BLA terminals in the NAcSh in control rats. **A.** Light delivery to BLA terminals in the NAcSh during deliberation in control rats (mCherry only) did not affect choice of the large, risky reward. **B.** Light delivery to BLA terminals in the NAcSh during delivery of the large, punished reward in control rats did not affect choice of the large, risky reward. Data are represented as mean ± SEM.

Additional analyses compared performance between eNpHR3.0 (n=13) and control (n=11) rats during the optogenetic test session. Sex was not included as a between-subjects factor as it was not a significant factor in the analyses of the data from each vector group separately. A three-factor repeated-measures ANOVA (session X vector group X trial block) revealed a main effect of session [*F* (1, 22) = 8,06, *p* = 0.01, ƞ^2^ = 0.25] as well as a significant session X trial block interaction [*F* (2, 44) = 5.28, *p* < 0.01, ƞ^2^ = 0.19]. Although there was neither a main effect of vector group [*F* (1, 22) = 0.02, *p* = 0.90, ƞ^2^ < 0.01] nor a session X vector group interaction [*F* (1, 22) = 0.99, *p* = 0.33, ƞ^2^ = 0.04], the session X vector group X trial group was significant [*F* (2, 44) = 1.43, *p* = 0.05, ƞ^2^ = 0.12]. This significant interaction confirms that, relative to baseline performance, light delivery to BLA terminals in the NAcSh increased risk taking only in eNpHR3.0 rats.

#### Inhibition during delivery of the large, punished reward

Because inhibition of BLA→NacSh during delivery of the large, punished reward affected risk taking specifically in eNpHR3.0 males, analysis of effects of light delivery during delivery of the large, punished reward was conducted only for control male rats (n=6). Light delivery during delivery of the large, punished reward did not affect choice of the large, risky reward [Figure 6B; session, *F* (1, 5) = 0.06, *p* = 0.82, ƞ^2^ = 0.01; session X trial block, *F* (2, 10) = 0.56, *p* = 0.59, ƞ^2^ = 0.10]. There were also no effects of light delivery on latencies to press levers during free choice trials [session, *F* (1, 5) = 3.25, *p* = 0.13, ƞ^2^ = 0.39; session X lever identity, *F* (1, 5) = 1.03, *p* = 0.36, ƞ^2^ = 0.17; session X lever identity X trial block, *F* (1, 5) = 0.57, *p* = 0.48, ƞ^2^ = 0.10]. Similarly, light delivery had no effect on latencies to collect food [session, *F* (1, 5) = 1.50, *p* = 0.28, ƞ^2^ = 0.23; session X trial block, *F* (1, 5) = 0.79, *p* = 0.41, ƞ^2^ = 0.14] nor did it impact the percentage of omitted free choice trials [Table 1; *t* (5) = −1.01, *p* = 0.36, d = 0.41]. Collectively, these data show that, similar to light delivery during deliberation, light delivery during the delivery of the large, punished reward also has no effect on risk taking in control male rats.

A three-factor repeated measures ANOVA was used to compare the effects of light delivery between eNpHR3.0 (n=8) and control male rats (n=6; this analysis was not conducted in females as there was no effect of inhibition in eNpHR3.0 females). Not only were there main effects of session [*F* (1, 12) = 18.14, *p* < 0.01, ƞ^2^ = 0.60] and vector group [*F* (1, 12) = 6.63, *p* = 0.02, ƞ^2^ = 0.36], but there were also significant session X vector group [*F* (1, 12) = 18.14, *p* < 0.01, ƞ^2^ = 0.63], session X trial block [*F* (2, 24) = 15.17, *p* < 0.01, ƞ^2^ = 0.56] and session X vector group X trial block [*F* (2, 24) = 23.02, *p* < 0.01, ƞ^2^ = 0.66] interactions. Results of these analyses confirm that the effects of light delivery during the delivery of the large, punished reward on risk taking is specific to eNpHR3.0 male rats.

### Experiment 2

#### In vitro electrophysiology

##### Green light inhibits D2R-expressing neurons in the NAcSh

Of the 52 D2-Cre transgenic rats used in Experiment 2, three (n=2, male; n=1, female) received intra-NAcSh infusions of the viral vector containing Cre-dependent eNpHR3.0 and were subsequently used for *in vitro* electrophysiology experiments to confirm that optogenetic inhibition of these neurons leads to a reduction in their activity. To directly evaluate light-induced inhibition of D2R-expressing neurons in the NAcSh of D2-Cre rats, cells expressing mCherry were targeted for analysis using whole-cell patch-clamp recordings (Figure 7A, B). On average, the membrane resistance and whole-cell capacitance of patched D2R-expressing neurons was 308.80 ± 39.67 MΩ and 62.59 ± 4.29 pF, respectively (n=32 cells). Exposure to green light reliably hyperpolarized cells and reduced or eliminated tonic firing (mean ΔV: −26.37 ± 3.67 mV; n=32, p=4.34^-8^, Figure 7C). Together, these findings demonstrate that green light robustly inhibits D2R-expressing neurons (i.e., those expressing Cre-dependent eNpHR3.0/mCherry) in the NAcSh.

**Figure 7.**
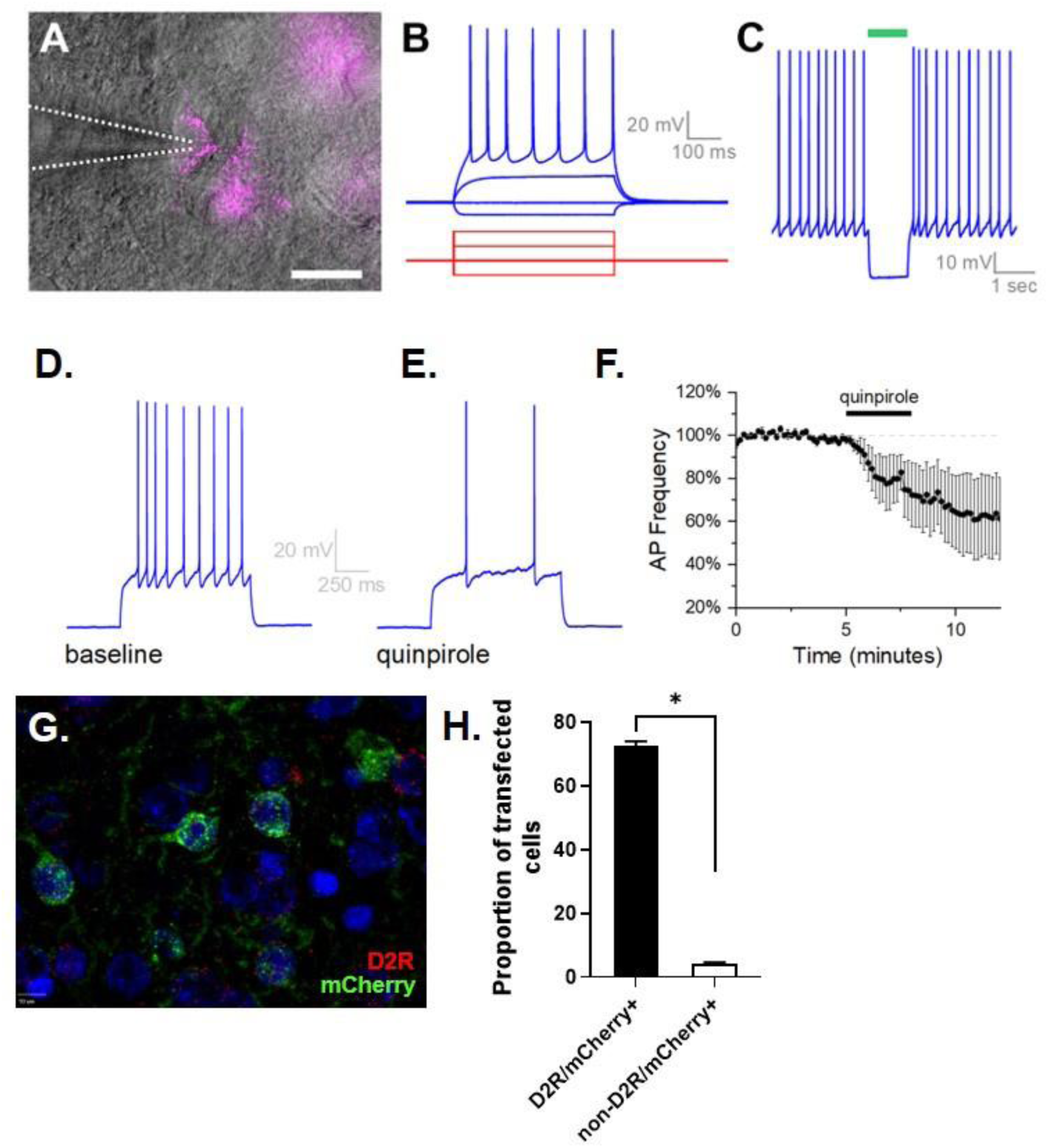
Functional and anatomical validation of the use of D2-Cre rats to optogenetically inhibit D2R-expressing neurons in the NAcSh. Green light inhibited eNpHR3.0/mCherry-positive D2R-expressing neurons in the NAcSh. A. NAcSh neurons expressing eNpHR3.0/mCherry were targeted by their red fluorescence (scale bar: 20 µm). B. Excitatory current delivered through the patch pipette induced action potential firing. C. Representative NAcSh neuron firing regularly in response to continuous 250 pA current injection is silenced by 1 sec green light exposure. D. Excitatory current injection (1 sec, 250 pA) was applied to induce continuous action potential firing, after which quinpirole was washed onto the tissue. E. Quinpirole (3 min, 10 µM) reduces the number of action potentials in response to the same current step. F. Normalized action potential frequency during the current step over time, as observed in all mCherry-expressing cells tested with quinpirole (n=18). G. Representative image depicting selective transduction of D2R-expressing neurons in the NAcSh. Green, mCherry; red, D2R mRNA; blue, DAPI. H. There were significantly more D2R/mCherry+ neurons than non-D2R/mCherry+ neurons in the NAcSh of rats injected with Cre-dependent eNpHR3.0-mCherry [t (23) = 38.36, p < 0.001, d = 7.83]. Data are represented as mean ± SEM. Asterisks indicate a significant difference (p < 0.05).

##### Quinpirole inhibits D2R-expressing neurons in the NAcSh

To evaluate the electrophysiological action of the D2R agonist quinpirole on activity of D2R-expressing neurons in the NAcSh, cells were targeted for study by their red fluorescence (i.e., mCherry). In neurons evaluated in voltage-clamp configuration at −70 mV, 3-minu53 exposure to 10 µM quinpirole produced an excitatory shift in holding current of 3.41 ± 1.4 pA and concomitantly reduced membrane resistance by 10.66 ± 2.30 % (n=10, *p* < 0.01). In neurons evaluated in current-clamp configuration, 3-minute exposure to 10 µM quinpirole reduced the frequency of action potentials observed during a suprathreshold depolarization by 39.1 ± 7.6% (n=18, *p* < 0.001, Figure 7D-F). Together, these findings provide evidence that eNpHR3.0/mCherry is expressed in D2R-expressing NAcSh neurons.

### Histology

A total of 52 D2-Cre transgenic rats (n=19, male; n=26, female) were used in Experiment 2. Of the 52 rats, 32 rats (n=19, female; n=13, male) received intra-NacSh infusions of the viral vector containing the Cre-dependent eNpHR3.0. Four (n=2/sex) of these rats were used for validation of selective viral transduction of D2R-expressing neurons using RNAscope (Figure 7G, H). Four females and two males in the eNpHR3.0 group lost their cranial implants prior to any optogenetic manipulations and were therefore removed from the study. Six females and one male were also excluded due lack of viral transduction (n=3), missed cannula placement (n=3), or infection around the site of the optic fiber (n=1). Nineteen rats (n=10, male; n=9, female) received intra-NAcSh infusions of the viral vector containing Cre-dependent mCherry. Of these rats, three females and one male lost their cranial implant prior to optogenetic manipulations and were therefore excluded from the study. Only one male was excluded due to a lack of viral expression in the NAcSh. After accounting for attrition, the final sample sizes were n=13 (n=7, male; n=6, female) for the eNpHR3.0 group and n=14 (n=7, male; n=7, female) for the mCherry group. Figure 7 depicts the locations of the tips of the optic fibers in the NAcSh in the eNpHR3.0 (Figure 8A) and control (Figure 8B) groups. As in Experiment 1, not all rats underwent every optogenetic test condition due to illness or loss of their cranial implant during the course of the experiment.

**Figure 8.**
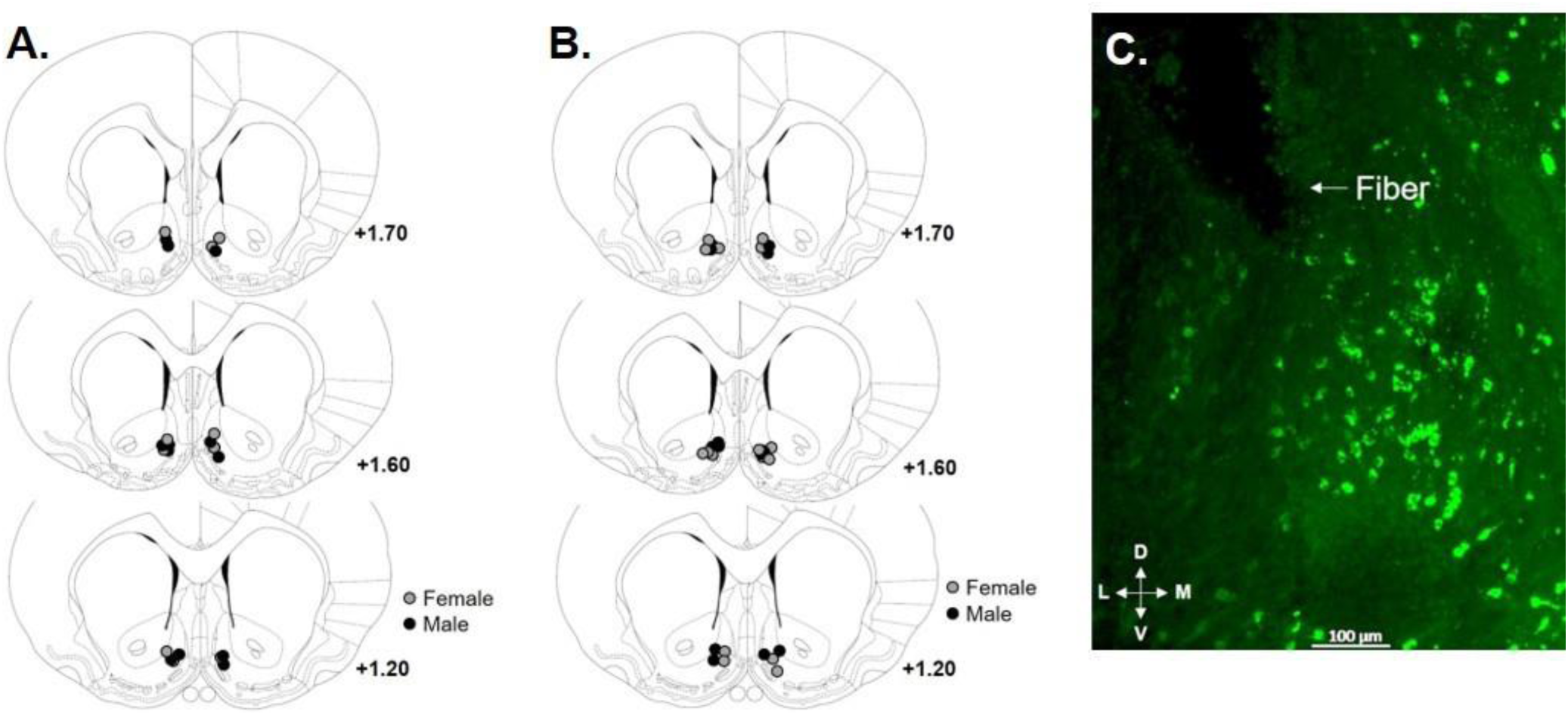
Optic fiber placement and eNpHR3.0 expression in the NAcSh. **A.** Circles represent the location of the tip of the optic fiber in the NAcSh of rats with expression of eNpHR3.0 in D2R-expressing neurons. **B.** Circles represent the location of the tip of the optic fiber in the NAcSh of rats with mCherry expression in D2R-expressing neurons. **C.** Representative image of the tip of the optic fiber in the NAcSh and expression of mCherry in D2R-expressing neurons in the NAcSh of an eNpHR3.0 rat.

### Optogenetic inhibition of D2R-expressing neurons in the NAcSh during decision making in eNpHR3.0 rats

#### Inhibition during deliberation

Inhibition of D2R-expressing neurons in the NAcSh during deliberation (n=5, male; n=6, female) increased choice of the large, risky reward [Figure 9A; session, *F* (1, 9) = 20.13, *p* < 0.01, ƞ^2^ = 0.69; session X sex, *F* (1, 9) = 0.15, *p* = 0.71, ƞ^2^ = 0.02; session X trial block, *F* (2, 18) = 11.15, *p* < 0.01, ƞ^2^ = 0.55; session X sex X trial block, *F* (2, 18) = 0.18, *p* = 0.83, ƞ^2^ = 0.02]. Analysis of the percentage of win-stay and lose-shift trials did not yield a main effect of session [*F* (1, 9) = 0.93, *p* = 0.36, ƞ^2^ = 0.09], but did reveal a significant session X trial type interaction [Figure 9B; *F* (1, 9) = 4.85, *p* = 0.05, ƞ^2^ = 0.35]. Similar to the effects of inhibition on percent choice of the large, risky reward, the effect of inhibition on trial type did not differ between males and females [session X sex, *F* (1, 9) = 0.65, *p* = 0.44, ƞ^2^ = 0.07; session X sex X trial type, *F* (1, 9) = 0.02, *p* = 0.90, ƞ^2^ < 0.01]. Consequently, data were collapsed across males and females, and *post-hoc* analyses were conducted to determine the source of the significant session X trial type interaction. While inhibition had no effect on the percentage of lose-shift trials relative to baseline [*t* (10) = 0.94, *p* = 0.37, d = 0.28], inhibition caused a significant increase in the percentage of win-stay trials [*t* (10) = −4.33, *p* < 0.01, d = 1.31]. The effects of inhibition on choice performance were specific to the optogenetic test session, as performance in the RDT 24 hours after the manipulation was no longer significantly different than baseline performance [session, *F* (1, 9) = 0.32, *p* = 0.59, ƞ^2^ = 0.03; session X sex, *F* (1, 9) = 0.71, *p* = 0.42, ƞ^2^ = 0.07; session X trial block, *F* (2, 18) = 0.01, *p* = 0.99, ƞ^2^ < 0.01; session X sex X trial block, *F* (2, 18) = 1.60, *p* = 0.23, ƞ^2^ = 0.15]. Together, these data reveal that activity of D2R-expressing neurons in the NAcSh during the deliberative period of decision making biases choice toward small, safe rewards, potentially via modulating reward sensitivity.

**Figure 9.**
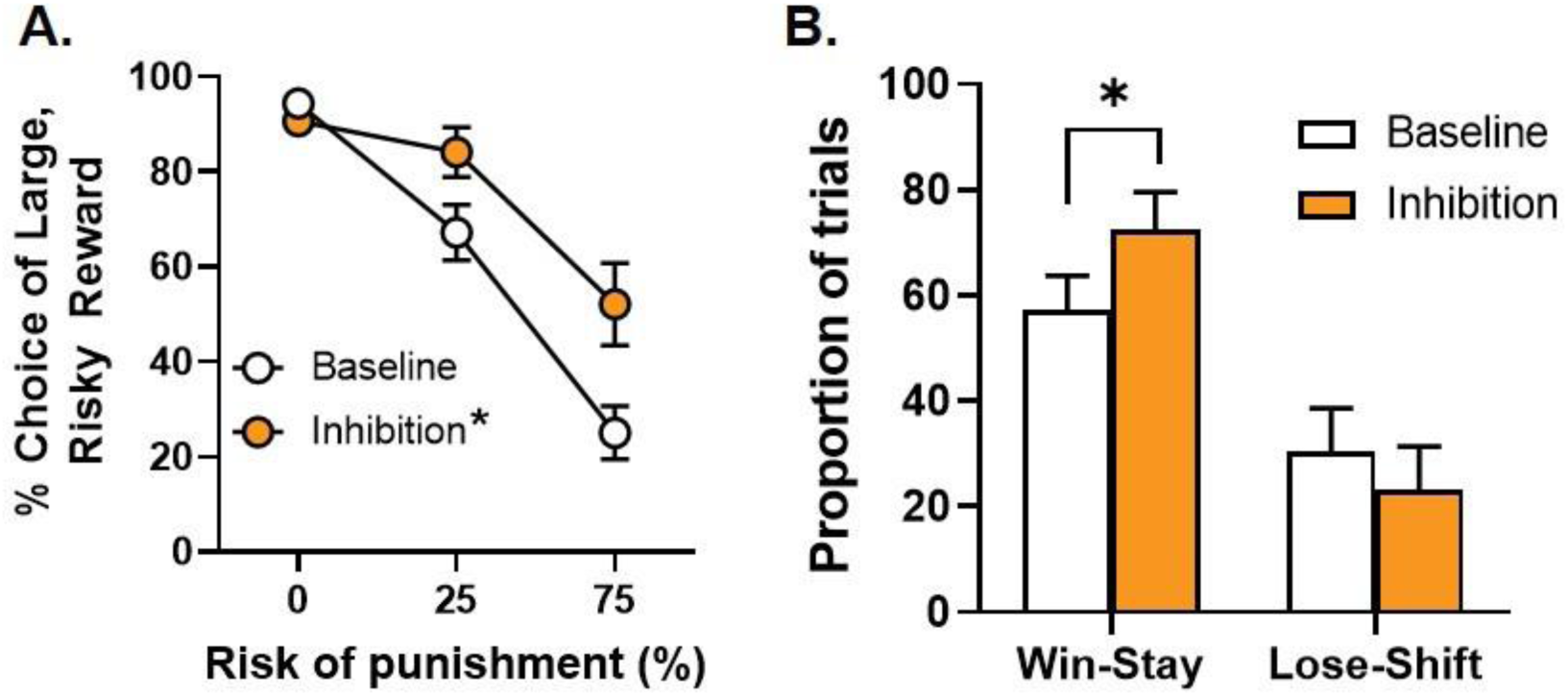
Optogenetic inhibition of D2R-expressing neurons in the NAcSh during deliberation. **A.** Inhibition of D2R-expressing neurons in the NAcSh during deliberation increased choice of the large, risky reward. **B.** Relative to baseline, inhibition of D2R-expressing neurons during deliberation increased the percentage of win-stay trials, but had no effect on the percentage of lose-shift trials. Data are represented as mean ± SEM. Asterisks indicate a significant difference between baseline (no laser delivery) and optogenetic inhibition (*p* < 0.05).

To determine whether inhibition of these neurons affected other behavioral measures in the RDT, latencies to nosepoke, latencies to lever press, and percentage of omissions were analyzed. Latencies to nosepoke to trigger lever extension were not affected by inhibition during deliberation [session, *F* (1, 9) = 2.01, *p* = 0.19, ƞ^2^ = 0.18; session X sex, *F* (1, 9) = 2.45, *p* = 0.15, ƞ^2^ = 0.21; session X trial block, *F* (2, 18) = 0.46, *p* = 0.64, ƞ^2^ = 0.05; session X sex X trial block, *F* (2, 18) = 0.14, *p* = 0.87, ƞ^2^ = 0.02]. Similar to latencies to nosepoke, there was no effect of inhibition on latencies to press levers during free choice trials [session, *F* (1, 6) = 1.72, *p* = 0.24, ƞ^2^ = 0.22; session X sex, *F* (1, 6) = 0.36, *p* = 0.57, ƞ^2^ = 0.06; session X lever identity, *F* (1, 6) = 1.11, *p* = 0.33, *p* = 0.16; session X lever identity X trial block, *F* (1, 6) = 0.56, *p* = 0.48, ƞ^2^ = 0.09; session X sex X trial block, *F* (1, 6) = 0.08, *p* = 0.79, ƞ^2^ = 0.01; session X lever identity X sex X trial block, *F* (1, 6) = 2.16, *p* = 0.19, ƞ^2^ = 0.26]. Finally, inhibition did not affect percentage of omitted free choice trials [Table 1; session, *F* (1, 9) = 0.04, *p* = 0.86, ƞ^2^ < 0.01; session X sex, *F* (1, 9) = 0.47, *p* = 0.51, ƞ^2^ = 0.05].

#### Inhibition during delivery of the small, safe reward

Optogenetic inhibition of D2R-expressing neurons in the NAcSh during delivery of the small, safe reward (n=6, male; n=5, female) caused a significant decrease in choice of the large, risky reward (Figure 10A). Although there was no main effect of session [*F* (1, 9) = 1.27, *p* = 0.07, ƞ^2^ = 0.32], there was a significant interaction between session and trial block [*F* (2, 18) = 9.35, *p* < 0.01, ƞ^2^ = 0.51]. This effect did not differ between males and females [session X sex, *F* (1, 9) = 0.18, *p* = 0.69, ƞ^2^ = 0.02; session X sex X trial block, *F* (2, 18) = 0.08, *p* = 0.93, ƞ^2^ < 0.01]. Inspection of Figure 10A shows that the significant interaction between session and trial block was mainly driven by an inhibition-induced decrease in choice of the large risky reward in the 25% trial block. Subsequent comparisons between baseline and optogenetic conditions for each block separately (correcting for multiple comparisons) confirmed that inhibition decreased risk taking only in the second block of trials [Blocks 1 and 3: session, *p-values* > 0.05, ƞ^2^s < 0.20; Block 2: session: *F* (1, 9) = 9.55, *p* = 0.01, ƞ^2^ = 0.52]. Analysis of the proportion of win-stay and lose-shift trials revealed no main effect of session [*F* (1, 6) = 2.14, *p* = 0.19, ƞ^2^ = 0.26] and no significant session X sex [*F* (1, 6) = 5.12, *p* = 0.07, ƞ^2^ = 0.46] or session X trial type [*F* (1, 6) = 0.20, *p* = 0.67, ƞ^2^ = 0.03] interactions. Despite these null effects, there was a significant session X sex X trial type interaction [*F* (1, 6) = 8.75, *p* = 0.03, ƞ^2^ = 0.59], suggesting that inhibition may have altered feedback processing differently between males and females. Sample sizes for each sex, however, were not sufficiently powered to perform *post-hoc* analyses within each sex separately to identify the source of this significant interaction (three males were not included in the analysis due to the lack of one or both of the trial types during the optogenetic session). Additional trial-by-trial analyses were subsequently conducted to determine the extent to which inhibition during the delivery of the small, safe reward affected subsequent choice. Trial types were categorized as “Stay” or “Shift”, with the former representing trials on which a rat chose the small, safe lever after receipt of a small reward (with optogenetic inhibition) and the latter representing trials on which a rat chose the large, risky lever after receipt of a small reward (also concurrent with inhibition). Because inhibition affected choice behavior specifically in the 25% risk block (Block 2), trial-by-trial analyses were conducted only on data from this block of trials. A three-factor repeated-measures ANOVA (session X trial type X sex) showed that although there was no main effect of session [*F* (1, 9) = 1.07, *p* = 0.33, ƞ^2^ = 0.11] nor significant session X sex [*F* (1, 9) = 0.77, *p* = 0. 40, ƞ^2^ = 0.08] or session X trial type X sex [*F* (1, 9) = 0.22, *p* = 0.65, ƞ^2^ = 0.02] interactions, there was a significant session X trial type interaction [Figure 10B; *F* (1, 9) = 10.63, *p* = 0.01, ƞ^2^ = 0.54]. *Post-hoc t-*tests (collapsed across sex) subsequently compared the proportion of each trial type between baseline and inhibition conditions. These analyses revealed that, relative to baseline, inhibition significantly increased the proportion of “Stay” trials [*t* (10) = −3.48, *p* < 0.01, d = 1.05], but significantly decreased the proportion of “Shift” trials [*t* (10) = 3.34, *p* < 0.01. d = 1.01]. These results indicate that when D2R-expressing neurons were inhibited during delivery of the small, safe reward, rats were more likely to continue to choose the lever associated with this reward rather than shifting to the lever associated with the more rewarding, yet riskier, option.

**Figure 10.**
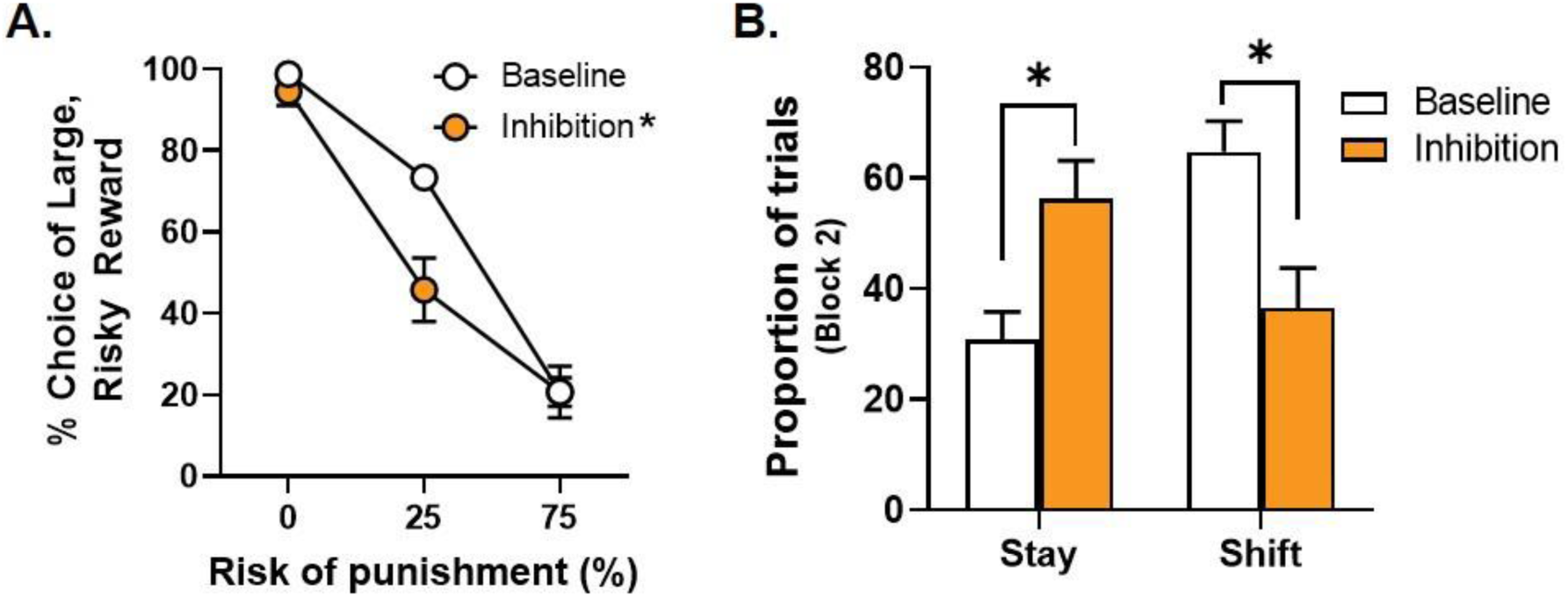
Optogenetic inhibition of D2R-expressing neurons in the NAcSh during delivery of the small, safe reward. **A.** Inhibition of D2R-expressing neurons in the NAcSh during the delivery of the small, safe reward decreased choice of the large, risky reward. **B.** Inhibition of D2R-expressing neurons during the delivery of the small, safe reward increased the percentage of “stay” trials and decreased the percentage of “shift” trials relative to baseline. Data are represented as mean ± SEM. Asterisks indicate a significant difference between baseline (no laser delivery) and optogenetic inhibition (*p* < 0.05).

Surprisingly, the effect of inhibition during delivery of the small, safe reward on risk taking persisted beyond the optogenetic test session. When baseline performance was compared to performance 24 hours after the optogenetic test session (collapsed across sex as there were no sex-dependent effects of inhibition on risk taking), there was a main effect of session [*F* (1, 10) = 9.10, *p* = 0.01, ƞ^2^ = 0.48], but no session X trial block interaction [*F* (2, 20) = 1.11, *p* = 0.35, ƞ^2^ = 0.10]. When baseline performance was compared to performance 48 hours after the optogenetic session, there was still evidence of the effects of inhibition on risk taking [session, *F* (1, 10) = 2.98, *p* = 0.12, ƞ^2^ = 0.23; session X trial block, *F* (2, 20) = 11.43, *p* < 0.01, ƞ^2^ = 0.53]. Together with the effects observed during the inhibition session itself, these results reveal that activity of D2R-expressing neurons in the NAcSh is important for the evaluation of less rewarding, albeit safer, options. Consequently, activity of these neurons during this evaluation period will bias choice toward more rewarding, yet costly, options. Further, disruption of activity of these neurons during this evaluation period may alter the way in which rats learn and encode reward contingencies in the task and therefore lead to longer term effects on risk-taking behavior (i.e., performance under subsequent non-inhibition conditions).

Comparison of latencies to press levers during free choice trials between baseline and the optogenetic test session yielded no main effect of session [*F* (1, 7) = 0.36, *p* = 0.57, ƞ^2^ = 0.05] nor a significant session X lever identity interaction [*F* (1, 7) = 0.32, *p* = 0.59, ƞ^2^ = 0.04]. There was a significant interaction between session, lever identity and trial block [*F* (1, 7) = 7.28, *p* = 0.03, ƞ^2^ = 0.51], but additional *post-hoc* analyses to determine the source of the significance did not reveal any main effects or significant interactions (all *p-values* > 0.05). Although females displayed overall longer latencies to press levers [*F* (1, 7) = 6.00, *p* = 0.04, ƞ^2^ = 0.46], there were no effects of inhibition on latencies to press levers in either males or females [session X sex, *F* (1, 7) < 0.01, *p* = 0.98, ƞ^2^ < 0.01; session X lever identity X sex, *F* (1, 7) = 0.12, *p* = 0.74, ƞ^2^ = 0.02; session X sex X trial block, *F* (1, 7) = 2.30, *p* = 0.17, ƞ^2^ = 0.25; session X lever identity X sex X trial block, *F* (1, 7) = 0.04, *p* = 0.84, ƞ^2^ < 0.01]. Similar to latencies to press levers, analyses of latencies to collect food were limited to data from Blocks 2 and 3 of the optogenetic test session due to rats’ near-exclusive preference for the large reward in Block 1. Inhibition during the delivery of the small, safe reward did not affect latencies to collect food [session, *F* (1, 9) = 2.43, *p* = 0.15, ƞ^2^ = 0.21; session X sex, *F* (1, 9) = 0.05, *p* = 0.83, ƞ^2^ < 0.01; session X trial block, *F* (1, 9) = 1.88, *p* = 0.20, ƞ^2^ = 0.17; session X sex X trial block, *F* (1, 9) = 3.76, *p* = 0.08, ƞ^2^ = 0.30]. Finally, there was no effect of inhibition on the percentage of omitted free choice trials [Table 1; session, *F* (1, 9) < 0.01, *p* = 1.00, ƞ^2^ < 0.01; session X sex, *F* (1, 9) = 0.10, *p* = 0.76, ƞ^2^ = 0.01].

#### Inhibition during delivery of the large, unpunished reward

Optogenetic inhibition of D2R-expressing neurons during delivery of the large, unpunished reward (n=5, male; n=5, female) did not affect choice of the large, risky reward [Figure 11A; session, *F* (1, 8) = 0.17, *p* = 0.69, ƞ^2^ = 0.02; session X sex, *F* (1, 8) = 0.01, *p* = 0.92, ƞ^2^ < 0.01; session X trial block, *F* (2, 16) = 1.19, *p* = 0.33, *p* = 0.13; session X sex X trial block, *F* (2, 16) = 0.15, *p* = 0.86, ƞ^2^ = 0.02]. Although females omitted significantly more free choice trials, [Table 1; *F* (1, 8) = 6.68, *p* = 0.03, ƞ^2^ = 0.46], there was no effect of inhibition on omissions in males or females [session, *F* (1, 8) < 0.01, *p* = 0.96, ƞ^2^ < 0.01; session X sex, *F* (1, 8) < 0.01, *p* = 0.97, ƞ^2^ < 0.01]. Hence, activity of D2R-expressing neurons in the NAcSh does not contribute to the evaluation of large, unpunished rewards during risk-taking behavior.

**Figure 11.**
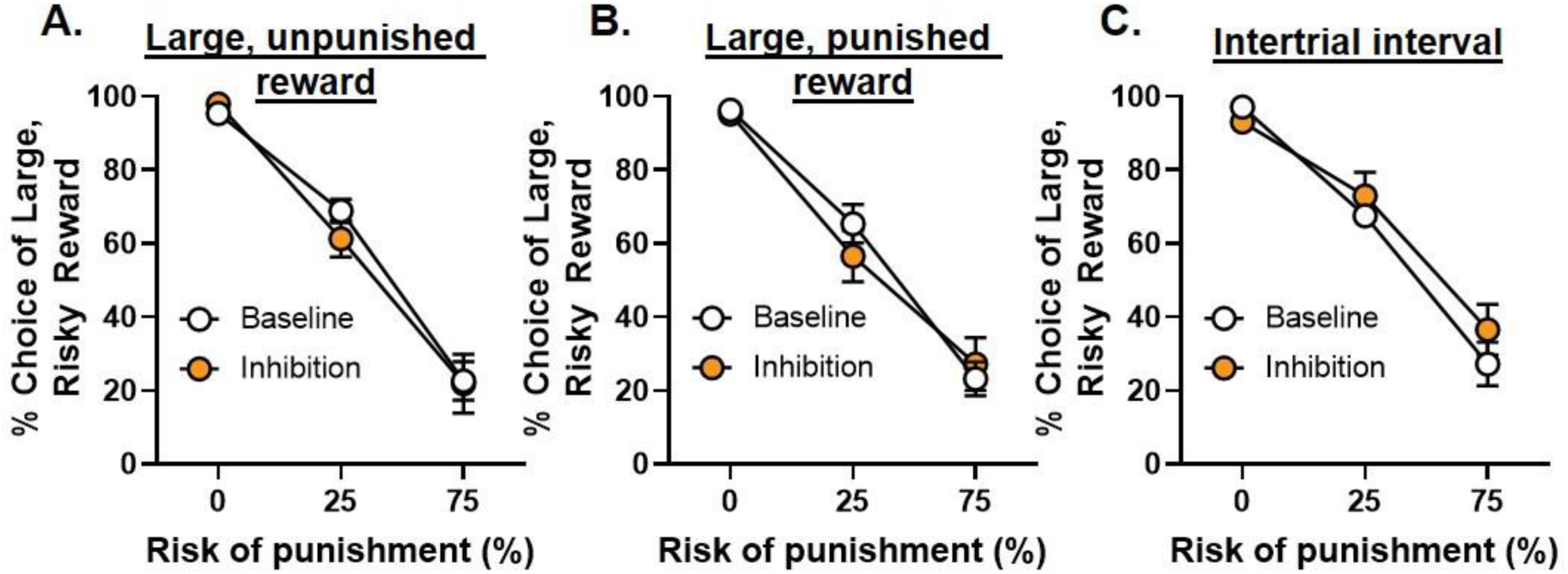
Optogenetic inhibition of D2R-expressing neurons in the NAcSh during other phases of the RDT. **A.** Inhibition of D2R-expressing neurons in the NAcSh during delivery of the large, unpunished reward did not affect choice of the large, risky reward. **B.** Inhibition of D2R-expressing neurons during delivery of the large, punished reward did not affect choice of the large, risky reward. **B.** Inhibition of D2R-expressing neurons during the intertrial interval did not affect choice of the large, risky reward. Data are represented as mean ± SEM.

#### Inhibition during delivery of the large, punished reward

Similarly, there were no effects of inhibition of D2R-expressing neurons during delivery of the large, punished reward (n=6, male; n=5, female) on choice of the large, risky reward [Figure 11B; session, *F* (1, 9) = 0.23, *p* = 0.64, ƞ^2^ = 0.03; session X sex, *F* (1, 9) < 0.01, *p* = 0.97, ƞ^2^ < 0.01; session X trial block, *F* (2, 18) = 1.30, *p* = 0.30, ƞ^2^ = 0.13; session X sex X trial block, *F* (2, 18) = 0.44, *p* = 0.65, ƞ^2^ = 0.05]. Consistent with previous results, females did omit significantly more free choice trials compared with males [Table 1; *F* (1, 9) = 7.90, *p* = 0.02, ƞ^2^ = 0.47], but inhibition did not affect omissions in either sex [*F* (1, 9) = 1.29, *p* = 0.29, ƞ^2^ = 0.13; session X sex, *F* (1, 9) = 1.90, *p* = 0.20, ƞ^2^ = 0.17]. These results suggest that, similar to the lack of involvement in the evaluation of unpunished rewards, activity of D2R-expressing neurons in the NAcSh does not appear to contribute to the evaluation of larger rewards that are accompanied by punishment.

#### Inhibition during the intertrial interval

As in Experiment 1, rats in Experiment 2 (n=4, male; n=6, female) also underwent an additional optogenetic test session in which light was delivered during the ITI. Not surprisingly, there was no effect of inhibition during the ITI on choice of the large, risky reward [Figure 11C; *F* (1, 8) = 1.19, *p* = 0.31, ƞ^2^ = 0.13; session X sex, *F* (1, 8) = 0.18, *p* = 0.68, ƞ^2^ = 0.02; session X trial block, *F* (2, 1. 16) = 2.12, *p* = 0.15, ƞ^2^ = 0.21; session X sex X trial block, *F* (2, 16) = 0.87, *p* = 0.44, ƞ^2^ = 0.10]. Additionally, there was no effect of inhibition during the ITI on the percentage of omitted free choice trials [Table 1; session, *F* (1, 8) < 0.01, *p* = 0.96, ƞ^2^ < 0.01; session X sex, *F* (1, 8) < 0.01, *p* = 0.98, ƞ^2^ < 0.01]. These results confirm that the inhibition-induced changes in risk taking during deliberation and delivery of the small, safe reward are not attributable to non-specific effects of eNpHR3.0 activation.

### Light delivery to D2R-expressing neurons in the NAcSh during decision making in control rats

Similar to Experiment 1, a separate cohort of transgenic rats (n=8, male; n=6, female) was used to confirm that the effects of inhibition of D2R-expressing neurons in the NAcSh were not due to light delivery alone. These rats were implanted with bilateral cannula in the NAcSh through which an AAV containing mCherry alone was infused and optic fibers were implanted. Like the eNpHR3.0 group, control rats were then trained in the RDT until they achieved stable baseline performance, after which optogenetic test sessions began. These sessions were limited to those in which inhibition of D2R-expressing neurons had significantly affected risk taking in the eNpHR3.0 group (i.e., deliberation and delivery of the small, safe reward). The order of these test sessions was counterbalanced across rats, and rats were required to re-acquire behavioral stability between successive optogenetic test sessions.

#### Inhibition during deliberation

In contrast to eNpHR3.0 rats, light delivery to D2R-expressing neurons in the NAcSh during deliberation (n=8, male; n=6, female) had no effect on choice of the large, risky reward in control rats [Figure 12A; session, *F* (1, 12) = 0.04, *p* = 0.85, ƞ^2^ < 0.01; session X sex, *F* (1, 12) = 0.72, *p* = 0.41, ƞ^2^ = 0.06; session X trial block, *F* (2, 24) = 0.14, *p* = 0.87, ƞ^2^ = 0.01; session X sex X trial block, *F* (2, 24) = 0.21, *p* = 0.81, ƞ^2^ = 0.02]. Light delivery did not affect latencies to nosepoke [session, *F* (1, 12) = 1.95, *p* = 0.19; session X sex, *F* (1, 12) = 0.08, *p* = 0.78; session X trial block, *F* (2, 24) = 2.63, *p* = 0.09, ƞ^2^ = 0.18; session X sex X trial block, *F* (2, 24) = 0.89, *p* = 0.42, ƞ^2^ = 0.07] nor did it alter latencies to press levers during free choice trials [session, *F* (1, 8) = 0.46, *p* = 0.52, ƞ^2^ = 0.06; session X sex, *F* (1, 8) = 0.12, *p* = 0.74, ƞ^2^ = 0.02; session X lever identity, *F* (1, 8) = 0.28, *p* = 0.61, ƞ^2^ = 0.03; session X sex X lever identity, *F* (1, 8) = 0.36, *p* = 0.57, ƞ^2^ = 0.04; session X lever identity X trial block, *F* (1, 8) = 0.45, *p* = 0.52, ƞ^2^ = 0.05; session X sex X lever identity X trial block, *F* (1, 8) = 0.52, *p* = 0.49, ƞ^2^ = 0.06]. Finally, there were no effects of light delivery on the percentage of omitted free choice trials [Table 1; session, *F* (1, 12) = 1.86, *p* = 0.20, ƞ^2^ = 0.13; session X sex, *F* (1, 12) = 1.21, *p* = 0.29, ƞ^2^ = 0.09. Together, these data indicate that light delivery in control rats had no effect on risk-taking behavior.

**Figure 12.**
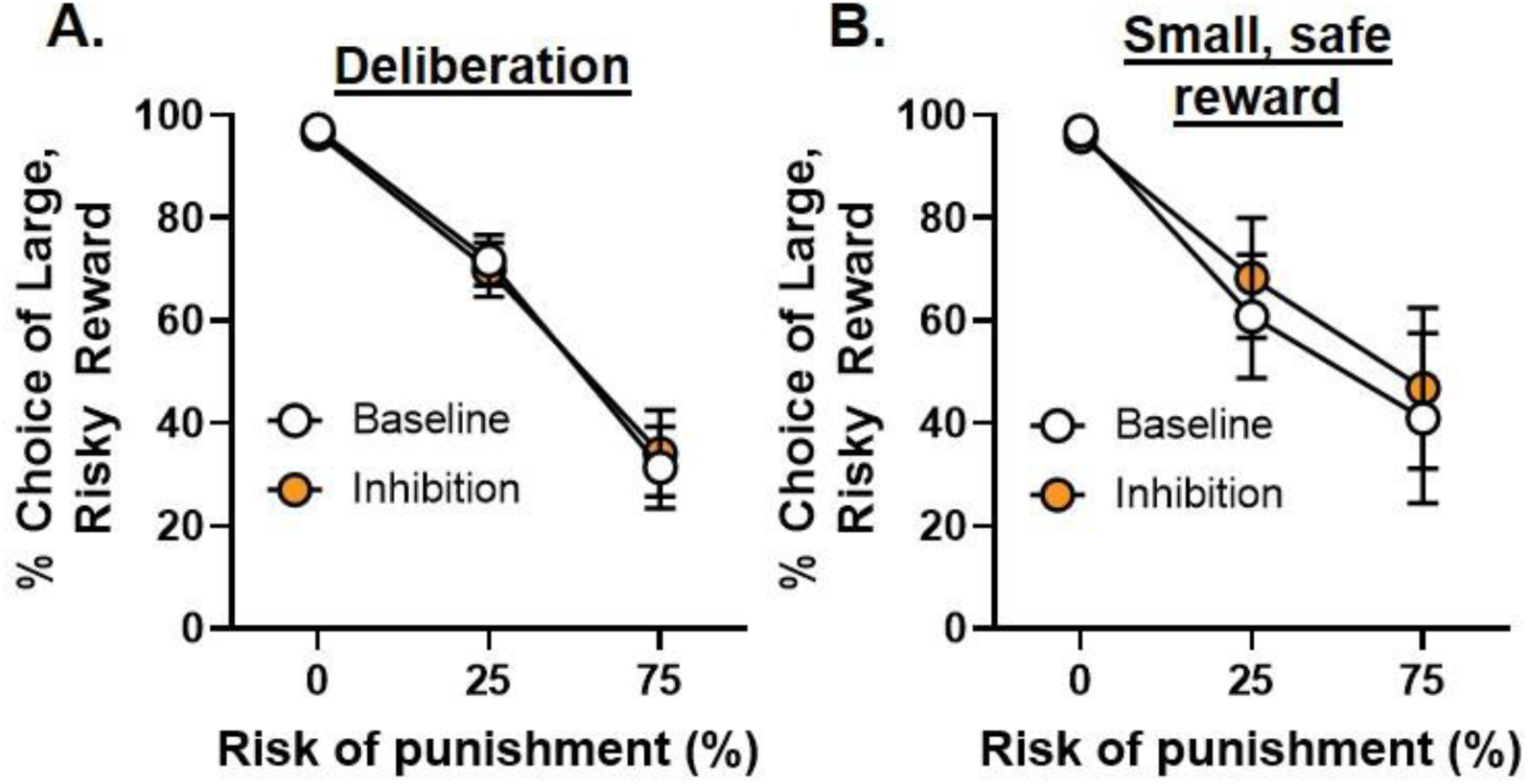
Light delivery to D2R-expressing neurons in the NAcSh in control rats. **A.** Light delivery to the NAcSh of control rats during deliberation did not affect choice of the large, risky reward. **B.** Light delivery to the NAcSh of control rats during the delivery of the small, safe reward did not affect choice of the large, risky reward. Data are represented as mean ± SEM.

As in Experiment 1, additional analyses were conducted to compare effects of light delivery during deliberation between eNpHR3.0 (n=11) and control rats (n=14). Because there were no sex-dependent effects of light delivery on risk taking in either group of rats, data were collapsed across sexes. Although there was no main effect of vector group [*F* (1, 23) = 0.20, *p* = 0.66, ƞ^2^ < 0.01], there was a main effect of session [*F* (1, 23) = 16.23, *p* < 0.01, ƞ^2^ = 0.41] and significant session X vector group [*F* (1, 23) = 13.24, *p* < 0.01, ƞ^2^ = 0.36], session X trial block [*F* (2, 46) = 8.05, *p* < 0.01, ƞ^2^ = 0.26] and session X vector group X trial block [*F* (2, 46) = 5.54, *p* < 0.01, ƞ^2^ = 0.19] interactions. Hence, these results confirm that, relative to baseline performance, light delivery during deliberation increased risk taking only in eNpHR3.0 rats.

#### Inhibition during delivery of the small, safe reward

Because of attrition, only two males remained in the control group that received intra-NacSh light delivery during delivery of the small, safe reward. Given this small sample size and the fact that there were no sex-dependent effects of inhibition in eNpHR3.0 rats, data were collapsed across sexes (n=2, male; n=4, female), yielding a sample size of n=6. Light delivery during delivery of the small, safe reward had no effect on choice of the large, risky reward [Figure 12B; session, *F* (1, 5) = 1.20, *p* = 0.23, ƞ^2^ = 0.19; session X trial block, *F* (2, 10) = 0.99, *p* = 0.41, ƞ^2^ = 0.17]. Light delivery did not alter latencies to press levers during free choice trials [session, *F* (1, 3) = 0.04, *p* = 0.85, 0.01; session X lever identity, *F* (1, 3) = 0.39, *p* = 0.58, ƞ^2^ = 0.12; session X trial block, *F* (1, 3) = 0.14, *p* = 0.74, ƞ^2^ = 0.04; session X lever identity X trial block, *F* (1, 3) = 0.02, *p* = 0.89, ƞ^2^ < 0.01] nor did it affect latencies to collect the small food reward [session, *F* (1, 4) = 3.59, *p* = 0.13, ƞ^2^ = 0.47; session X trial block, *F* (1, 4) = 0.02, *p* = 0.89, ƞ^2^ < 0.01]. There was also no effect of light delivery on the percentage of omitted free choice trials [Table 1; *t* (10) = −0.03, *p* = 0.97, d = 0.01].

A comparison of the effects of light delivery between eNpHR3.0 (n=11) and control rats (n=6; collapsed across sex) revealed that although there were no main effects of session [*F* (1, 15) = 0.85, *p* = 0.37, ƞ^2^ = 0.05] or vector group [*F* (1, 15) = 1.41, *p* = 0.25, ƞ^2^ = 0.09], there were significant session X vector group [*F* (1, 15) = 4.23, *p* = 0.05, ƞ^2^ = 0.22], session X trial block [*F* (2, 30) = 3.31, *p* = 0.05, ƞ^2^ = 0.18] and session X vector group X trial block [*F* (2, 30) = 6.13, *p* < 0.01, ƞ^2^ = 0.29] interactions. These results confirm that the effects of light delivery during the delivery of the small, safe reward were specific to rats in the eNpHR3.0 group.

## Discussion

By leveraging the temporal precision afforded by optogenetics, the current study dissected the roles of the BLA→NAcSh (Experiment 1) and neurons that selectively express D2Rs in the NAcSh (Experiment 2) in different phases of decision making involving risk of punishment. Inhibition of the BLA→NAcSh during either deliberation or delivery of the large, punished reward increased risk taking, although the latter effect was observed only in males. In contrast, inhibition of NAcSh D2R-expressing neurons had distinct effects depending on the phase of the decision process. Whereas inhibition of these neurons during deliberation increased risk taking, inhibition during delivery of the small, safe reward decreased risk taking. These findings advance our understanding of the neurobiology of risk-based decision making by revealing the importance of the BLA→NAcSh in processing risk-related information and the dissociable role of NAcSh D2R-expressing neurons in guiding risk-taking behavior.

### The role of the BLA→NAcSh in risk-based decision making

Recently, we demonstrated that the BLA is differentially recruited during decision making involving risk of punishment (Orsini et al., 2017), in that inhibition during deliberation decreased risk taking, whereas inhibition during delivery of the large, punished reward increased risk taking. We hypothesized these dissociable effects may be due to engagement of distinct BLA cell populations with divergent projections to downstream brain regions during different phases of the decision process. For example, based on prior work showing a temporally-specific role for the BLA→NAc in decision making involving risk of reward omission (e.g., probability discounting task; Bercovici et al., 2018), BLA→NAcSh may be particularly important for evaluation of punished rewards to provide negative feedback that guides future choices. Consistent with this hypothesis, inhibition of BLA→NAcSh during delivery of the large, punished reward increased risk taking in the RDT. Surprisingly, however, this effect was only observed in males, with inhibition failing to alter risk taking in females. Such an effect in males is consistent with our previous work showing increased risk taking in male rats when BLA inhibition occurred during the same phase of decision making (Orsini et al., 2017). Further, this inhibition-induced increase in risk taking is congruent with the increase in risky choice observed when the BLA→NAc is inhibited following a non-rewarded risky choice in the probability discounting task (Bercovici et al., 2018). Collectively, these studies suggest the BLA→NAc is important for evaluation of negative outcomes (either punishment or lost reward opportunity) that can be used to guide future choices toward safer or more certain rewards.

The observation that BLA→NAcSh inhibition during delivery of the large, punished reward had no effect in females suggests the neural mechanisms by which males and females evaluate punished rewards are different. Previous work in both humans and rodents demonstrates that BLA activity is greater in females compared with males at baseline and in response to a potentially aversive stimulus (Lebron-Milad et al., 2012; Cover et al., 2014; Blume et al., 2017; Keiser et al., 2017; Blume et al., 2019; Hodges et al., 2022). Similarly, a c-fos mapping study revealed greater NAcSh activity and, more notably, greater functional connectivity between the BLA and NAc in females compared with males (Hodges et al., 2022). Hence, rather than a lack of BLA→NAcSh involvement in evaluation of negative feedback during risk taking in females, BLA→NAcSh activity may instead be greater in females than males during this decision-making phase. Consequently, optogenetic inhibition using our current parameters, which were based on prior work in which BLA→NAcSh inhibition effectively altered motivated behavior (Stuber et al., 2011), may not have been sufficient to disrupt this process in females. Alternatively, the absence of an effect in females could be due to anatomical differences, such as fewer projections BLA →NAcSh projections in females relative to males (although to our knowledge there is no evidence to support such differences). Finally, though it is possible that differences in shock intensities used for males and females could have contributed to the divergent effects of BLA-NAcSh inhibition on risk taking, this explanation is also unlikely as shock intensities were titrated individually for each rat such that their choice performance was neither at the floor nor at the ceiling of the parametric space. In doing so, shock intensities ranged across the sexes, with some males tested with lower shock intensities than females.

Contrary to our initial hypothesis, there was a modest, but significant, increase in risk taking following BLA→NAcSh inhibition during deliberation, which was evident in both males and females. These results were surprising for two reasons. First, optogenetic inhibition of BLA glutamatergic neurons during deliberation decreases, rather than increases, risk taking (Orsini et al., 2017). Second, BLA→NAc inhibition during deliberation decreases choice of a large, risky reward when the risk (in this case, risk of reward omission) is low (Bercovici et al., 2018). Based on these studies, one would therefore predict that inhibition would decrease, rather than increase, risk taking, particularly in the 25% risk block. This discrepancy can be reconciled, however, after considering the target of inhibition and the structure of the decision process itself. In contrast to inhibition of all BLA glutamatergic neurons, which would silence numerous BLA efferent projections, inhibition of a specific projection from the BLA may result in novel and distinct changes in behavior. More broadly, decision making is not a linear process and thus, information that is processed in one phase of a decision may be integrated in other phases. Accordingly, BLA→NAcSh inhibition during deliberation might prevent the integration of negative feedback information (from the previous evaluation phase) into the deliberative process, resulting in effects on risk taking similar to those seen when inhibition occurred during delivery of the large, punished reward. These findings also suggest that the decrease in risk taking following BLA inhibition during deliberation (Orsini et al., 2017) is not due to inhibition of BLA neurons that project to the NAcSh. Rather, the BLA must communicate with separate downstream target(s), such as the orbitofrontal cortex (Orsini et al., 2015a), during the deliberative process to bias choices toward larger, riskier options.

#### The role of NAcSh D2R-expressing neurons in risk-based decision making

In contrast to other forms of risk-based decision making (St Onge and Floresco, 2009; Stopper et al., 2013; Di Ciano et al., 2015), D2Rs have a selective role in decision making involving risk of punishment (Simon et al., 2011; Blaes et al., 2018). In particular, intra-NAc infusion of the D2R agonist quinpirole reduces risk taking, indicating a role for D2Rs in this brain region (Mitchell et al., 2014). These studies are limited, however, by their inability to dissect the role of D2Rs in a temporally-specific manner. The current study circumvented this limitation by using optogenetics in a transgenic rat line that allows selective manipulation of D2R-expressing neurons. These results supported the premise that optogenetic inhibition of these neurons may be a tractable approach to mimic effects of D2R agonist administration, but in a more temporally-precise manner.

When inhibition of these neurons occurred during deliberation, there was an increase in risk taking. These findings are consistent with a recent study in which optogenetic excitation of NAc D2R-expressing neurons during deliberation decreased risky choice (i.e., choice of a reward that varied in its magnitude and frequency of delivery; Zalocusky et al., 2016). Together with the current study, these findings suggest activity of D2R-expressing neurons during deliberation promotes choice away from large, risky rewards. In the current study, however, inhibition-induced increases in risk taking were accompanied by selective increases in the percentage of win-stay trials, indicating that the increase in risk taking was driven by an augmented sensitivity to the larger, more rewarding outcome. In contrast, Zalocusky et al. (2016) attributed their effects to alterations in sensitivity to losses. This discrepancy may be due to the difference in the nature of the risks associated with the “risky” option: footshock in the current study versus reward omission in Zalocusky et al. (2016). Indeed, D2R agonists reduce risk taking under risk of punishment, whereas they increase risk taking under risk of reward omission (St Onge and Floresco, 2009; Simon et al., 2011). Hence, the contribution of these neurons to the integration of feedback information during deliberation may differ depending on the nature of the risk associated with the available options during decision making.

Notably, the effect of inhibition of D2R-expressing neurons during deliberation was similar to effects of BLA→NAcSh inhibition on risk taking, suggesting BLA neurons may communicate directly with NAcSh D2R-expressing neurons during deliberation to modulate choice behavior. Indeed, BLA excitatory projections to the NAcSh synapse onto D2R-expressing medium spiny neurons (MSNs), although they also synapse onto D1R-expressing MSNs (MacAskill et al., 2014; Barrientos et al., 2018; Zinsmaier et al., 2022). Anatomical and physiological evidence suggests, however, that BLA excitatory projections regulate NAc activity through modulation of D1R-, but not D2R-, expressing MSNs (Floresco et al., 2001; Charara and Grace, 2003). Although the BLA could influence D2R-expressing neurons indirectly via intermediary D1R MSNs, recent work has shown that BLA excitatory projections to the NAcSh predominantly synapse onto inhibitory fast-spiking interneurons (FSIs), and activation of FSIs by BLA input precedes activation of MSNs (Yu et al., 2017). Given that D1Rs are not involved in risk-taking behavior (Simon et al., 2011), it seems more likely that BLA activity during deliberation influences D2R-expressing MSNs through direct activation of intermediary FSIs. It is also possible that similar effects of inhibition during deliberation across the BLA→NAcSh and D2R-expressing neurons is coincidental, and that activity of D2R-expressing neurons is regulated by alternate afferent regions, such as the ventral tegmental area (Gerfen et al., 1987; Heimer et al., 1997). Future experiments will distinguish between these (and other) potential explanations for how activity of D2R-expressing neurons is regulated during deliberation to guide subsequent risky choice.

In contrast to the effects of inhibition during deliberation, inhibition during delivery of the small, safe reward decreased risk taking. Trial-by-trial analysis revealed that rats were more likely to continue to choose the small, safe option again after concurrent inhibition and small reward delivery. Such a decrease in risk taking is congruent with effects of intra-NAc infusions of quinpirole on risk taking (Mitchell et al., 2014), suggesting that this pharmacological manipulation may exert its effects primarily on this reward evaluation period. The current results suggest that activity in these neurons during this decision-making phase is important for biasing choice toward more rewarding options. This interpretation is consistent with recent work showing that optogenetic activation of NAc D2R-expressing MSNs enhances motivation for food (Soares-Cunha et al., 2016; Soares-Cunha et al., 2018). If activity of these neurons enhances reward motivation during risk taking, inhibition should have also decreased risk taking during delivery of the large reward; however, inhibition was ineffective during this phase. Alternatively, activity in D2R-expressing neurons during evaluation of the small, safe reward may be important for balancing the relative reward of the alternate option against its associated risk. When the risk is low (as in the 25% trial block), the risk is perceived as minimal and therefore it may be more adaptive to shift choice toward the larger reward. When these neurons are inhibited, the risk overshadows the value of the larger reward, resulting in continued choice of the safer, albeit less rewarding, option.

Although the D2-Cre transgenic line allowed selective manipulation of D2R-expressing neurons, it did not allow differentiation among different striatal cell types expressing D2Rs, which are present on both GABAergic MSNs and cholinergic interneurons (CINs; Nicola et al., 2000; Kreitzer, 2009; Simpson et al., 2022). Although CINs constitute only ∼5% of NAc neurons, they exert strong regulatory influence over other populations, including MSNs. It will therefore be important to continue to develop tools to allow further cell-specific dissection of the role of D2Rs in different aspects of decision making. One intriguing hypothesis is that the dissociable effects of inhibiting D2R-expressing neurons during different decision phases is due to inhibition predominantly affecting one D2R-expressing cell type over the other.

## Conclusions

The current findings advance our understanding of the neural mechanisms that govern distinct aspects of risk-based decision making. They also provide evidence that circuits involved in decision making may be recruited in a sex-dependent manner during distinct phases of the decision process. In addition, the observation that selective cell populations are differentially engaged during decision making reaffirms the importance of using temporally-specific manipulations to fully understand the neural substrates inherent in such complex cognitive operations. Finally, our findings provide insight into the neural mechanisms by which risk-based decision making may become compromised in psychiatric diseases and suggest approaches to mitigate these cognitive impairments.

## Acknowledgements

Acknowledgements

We thank Ms. Bonnie McLaurin and Ms. Merrick Garner for their technical assistance. Supported by, F31DA057112-01 and a Bruce/Jones Graduate Fellowship (LMT), R00DA041493 (CAO), RF1AG060778 (JLB, BS and CJF).

